# mTORC1 Regulates the Metabolic Switch of Postnatal Cardiomyocytes During Regeneration

**DOI:** 10.1101/2023.09.12.557400

**Authors:** Wyatt G. Paltzer, Timothy J. Aballo, Jiyoung Bae, Katharine A. Hubert, Dakota J. Nuttall, Cassidy Perry, Kayla N. Wanless, Raya Nahlawi, Ying Ge, Ahmed I. Mahmoud

## Abstract

The metabolic switch from glycolysis to fatty acid oxidation in postnatal cardiomyocytes contributes to the loss of the cardiac regenerative potential of the mammalian heart. However, the mechanisms that regulate this metabolic switch remain unclear. The protein kinase complex mechanistic target of rapamycin complex 1 (mTORC1) is a central signaling hub that regulates cellular metabolism and protein synthesis, yet its role during mammalian heart regeneration and postnatal metabolic maturation is undefined. Here, we use immunoblotting, rapamycin treatment, myocardial infarction, and global proteomics to define the role of mTORC1 in postnatal heart development and regeneration. Our results demonstrate that the activity of mTORC1 is dynamically regulated between the regenerating and the non-regenerating hearts. Acute inhibition of mTORC1 by rapamycin or everolimus reduces cardiomyocyte proliferation and inhibits neonatal heart regeneration following injury. Our quantitative proteomic analysis demonstrates that transient inhibition of mTORC1 during neonatal heart injury did not reduce protein synthesis, but rather shifts the cardiac proteome of the neonatal injured heart from glycolysis towards fatty acid oxidation. This indicates that mTORC1 inhibition following injury accelerates the postnatal metabolic switch, which promotes metabolic maturation and impedes cardiomyocyte proliferation and heart regeneration. Taken together, our results define an important role for mTORC1 in regulating postnatal cardiac metabolism and may represent a novel target to modulate cardiac metabolism and promote heart regeneration.

## INTRODUCTION

The inability of the adult mammalian heart to regenerate damaged tissue after injury has resulted in a significant health and economic burden from widespread heart failure with reduced ejection fraction [1]. In contrast, the neonatal mouse heart possesses a remarkable ability to regenerate after injury, in stark contrast to the adult mouse heart [2, 3]. Shortly after birth, cardiomyocytes transition from hyperplastic to hypertrophic growth, resulting in loss of the cardiac regenerative potential [4-6]. One key biological process of postnatal cardiomyocyte maturation is a switch in metabolic substrate utilization. Highly proliferative embryonic and neonatal cardiomyocytes rely upon glycolysis, but exposure to an oxygen-rich environment after birth stimulates a metabolic switch towards the more energy efficient mitochondrial oxidative phosphorylation and fatty acid utilization as the primary energy source [5-7]. The cardiac metabolic state can have important implications in modulating cardiac maturation and regeneration. However, the mechanisms that control this metabolic switch in postnatal cardiomyocytes is not fully understood.

mTORC1 has been implicated as a master regulator of protein synthesis and cellular metabolism in response to nutrient and growth signals, where mTORC1 function regulates the levels of key metabolic enzymes involved in glycolysis, oxidative phosphorylation, and fatty acid synthesis [8]. mTORC1’s regulation of protein synthesis and metabolism is mediated via direct phosphorylation of its primary downstream targets eukaryotic translation initiation Factor 4E binding protein 1 (4EBP1) and ribosomal protein S6 kinase (S6K). mTORC1 promotes glycolysis through phosphorylation of 4EBP1 and S6K, which increases levels of master regulators of glycolysis such as hypoxia-inducible factor 1α (HIF1α) and MYC that can induce the expression of glycolytic enzymes [9-11]. In addition, mTORC1 controls oxidative phosphorylation through multiple regulatory mechanisms at the transcriptional and translational level [12, 13]. However, whether mTORC1 regulates the postnatal metabolic switch in cardiomyocytes remains undetermined.

mTOR plays an essential role during cardiac development and homeostasis. mTOR inhibition through cardiomyocyte-specific deletion of Mtor leads to lethality during both embryonic and postnatal development due to impaired cardiomyocyte survival and proliferation [14, 15]. Furthermore, inhibition of mTORC1 disrupts adult cardiac function, where inducible cardiomyocyte-specific knockout of the mTORC1 subunit; Raptor; is lethal within 8 weeks of due to increased apoptosis and disruption of metabolic substrate use [16]. Interestingly, recent studies demonstrate that modulating mTORC1 function through modifications to the upstream regulator tuberous sclerosis complex (TSC2) can regulate cardiac metabolism following ischemia reperfusion injury [17, 18]. These results demonstrate an important role for mTORC1 in postnatal heart function and metabolism.

mTORC1 is a critical node in regulating key metabolic pathways; however, the role of mTORC1 in the postnatal metabolic switch, cardiomyocyte cell cycle activity, and cardiac regeneration remains unclear. In this study, we define the distinct function of mTORC1 in the postnatal heart as well as following injury. We demonstrate that transient inhibition of mTORC1 activity blocks cardiomyocyte proliferation and neonatal mouse heart regeneration. Furthermore, our quantitative proteomic analysis demonstrates that inhibition of mTORC1 in the postnatal heart induces an accelerated metabolic switch from glycolysis to fatty acid oxidation. Our results reveal a novel role for mTORC1 function in regulating postnatal cardiomyocyte metabolism and mammalian cardiac regenerative potential.

## METHODS

### Animals

Untimed pregnant CD1 females were obtained from Charles River Laboratories, and neonatal mice were then used for all experiments. All experimental procedures involving animals were approved by the Institutional Animal Care and Use Committee of the University of Wisconsin-Madison. All animal experiments were performed with age-matched mice.

### Myocardial Infarction (MI) Model

Neonatal mice at postnatal day (P) 1 underwent a surgically induced MI as described previously [3]. Neonates were anesthetized via hypothermia by placing them on ice for up to 5 min. Once anesthetized, a lateral incision was made into the skin to view the muscle above the fourth intercostal space. A lateral thoracotomy was then performed by blunt dissection of the fourth intercostal space. A C-1 tapered needle connected to 6-0 prolene suture (Ethicon Inc, Bridgewater, NJ) was then passed through the ventricle around the left anterior descending coronary artery, which was then tied off to induce a clinically significant MI. The lateral thoracotomy was then closed by using the prolene suture to tie the ribs together, whereas the skin incision was closed by using surgical adhesive glue (3M). Mice were then warmed in-hand until they began to awaken, mice were then placed onto a heating pad until fully recovered. Sham-operated mice underwent the same procedure, including anesthetization, however their coronary artery was not ligated.

### Drug Administration

Neonatal mice were weighed daily and injected with saline, 1.0 mg/kg Rapamycin, or 1.0 mg/kg Everolimus. Stock solutions of both Rapamycin and Everolimus were diluted 100 mg/ml in dimethyl sulfoxide (Sigma) and then further diluted in saline prior to injection. Saline was used as the vehicle control for all experiments. All treated mice were given intraperitoneal injections on days 4, 5, and 6 post-MI. In mice treated with puromycin for measuring protein translation, mice were weighed 1 hr prior to harvest and injected with 0.04 µmol/g puromycin (VWR, #75844-852).

### Immunoblotting Assay

Myocardial ventricular lysates were obtained from ventricular tissue flash frozen on liquid nitrogen and prepared in RIPA buffer (Thermo Fisher, #89900) with 1X HALT protease and phosphatase inhibitor cocktail (Thermo Fisher, #78440). Protein concentration for each sample was then determined by BCA assay (Thermo Fisher, #23225). Samples were then prepared in SDS-PAGE loading buffer (LI-COR, #928-40004) and ran on either 10% or 15% polyacrylamide containing gels with transfer blotting on to a polyvinylidene fluoride membrane. Proteins of interest were targeted with the following antibodies, phospho-p70 S6K T389 (CST, #9205, 1:1000), p70 S6K (CST, #9202, 1:1000), phospho-4EBP1 T37/T46 (CST, 1:1000), phospho-4EBP1 S65 (CST, #9451, 1:1000), 4EBP1 (CST, 9452, 1:1000), puromycin (Millipore Sigma, #MABE343, 1:5000), and GAPDH (Proteintech, 60004-1, 1:1000). The antibody binding was then visualized using the Odyssey FC (LI-COR) imaging system and Image Studio Software version 4.0. Staining quantification was then conducted using the Image Studio Software version 4.0.

### Immunofluorescent Staining

Paraffin embedded hearts were sectioned and placed onto glass slides, slides were then deparaffinized, rehydrated, and boiled in IHC antigen retrieval solution (Invitrogen, #00-4955- 58). Sections were then blocked with 5% goat serum (Vector Laboratories, #S-1000) and incubated overnight at 4°C with primary antibodies. Primary antibodies used in this study were phospho-histone H3 Ser10 (Millipore, catalog # 06-570, 1:100), aurora B Kinase (Sigma, catalog # A5102, 1:100), and cardiac troponin T (Abcam, catalog # ab829, 1:100). Sections were washed and incubated with a corresponding secondary antibody conjugated to Rabbit-488 (Invitrogen, #A-11008, 1:400) or Mouse-555 (Invitrogen, #A28180, 1:400) for 1 hr at room temperature, with nuclei then stained using DAPI (Sigma, #D9542). Slides were mounted using Fluoromount-G (Thermo Fisher, #00-4958-02) and imaged for quantification on a Keyence BZ-X800 microscope, with high magnification images taken on a Nikon A1RS confocal microscope.

### Trichrome Staining

Hearts were harvested and fixed using 4% paraformaldehyde (PFA) in PBS overnight at 4 °C. The tissue was then washed with PBS and incubated overnight in 70% ethanol for dehydration before processing and paraffin embedding. Cardiac fibrosis and scar formation was then assayed using Masson’s trichrome staining with Newcomer Supply’s “Trichrome, Masson, Aniline Blue Stain Kit” (Newcomer Supply, #9179B) following the kit protocol. At least 4 biological replicates were analyzed with at least 3 independent sections from each biological replicate being stained for analysis.

### Cardiomyocyte Cross-Sectional Area Quantification

To quantify cardiomyocyte cross sectional area, we stained paraffin embedded heart sections with a wheat germ agglutinin (WGA) (Thermo Fisher, #W11261) antibody that was pre-conjugated to an Alexa Fluor 488 fluorophore for visualization. Slides were incubated with the WGA antibody for 1 hr at room temperature, rinsed with PBS-Tween (0.5%) and mounted in Fluoromount-G (Thermo Fisher, #00-4958-02) mounting medium, after which slides were then imaged. Fiji was used to quantify cardiomyocytes by measuring the cross-sectional area of 150 cardiomyocytes across 5 biological replicates for MI and 4 biological replicates for sham (SH) with 3 sections measured per biological replicate.

### Global Quantitative Proteomic Analysis

Hearts were excised from the mice, the atria were dissected from the heart, and the hearts were quickly washed with PBS. Ventricular tissue was then snap-frozen with liquid nitrogen and cryopulverized. The global ventricular proteome was extracted as previously described with Azo lysis buffer with minor modification [19, 20]. In a cold room (4 LJ), ventricular tissue was thoroughly homogenized in 0.2% Azo lysis buffer (0.2% w/v Azo, 25 mM ammonium bicarbonate (ABC), 10 mM L-methionine, 1 mM DTT, and 1X HALT protease and phosphatase inhibitor) with a Teflon pestle. Samples were then sonicated in a sonicating water bath for 10 min at 4 □ prior to centrifugation at 21,100 *g* for 30 min at 4 LJ. The supernatant was transferred to a new tube, and sample aliquots were diluted 1:50 in water before Bradford protein assay (Bio-Rad, Hercules, CA, USA, Cat# 5000006). Samples were normalized to 1 mg/mL in 0.1% Azo, reduced with 30 mM DTT at 37 □ for 1 h and alkylated with 30 mM chloroacetamide for 45 min. Trypsin Gold (Promega, Madison, WI, USA, Cat# V5820) was added to each sample in a 1:50 ratio (wt/wt) of protease:protein and incubated on 1000 rpm shaker at 37 □ overnight. The reaction was halted with 1% FA (final concentration). Azo was degraded at 305 nm (UVN-57 Handheld UV Lamp; Analytik Jena, Jena, TH, DEU) for 5 min, samples were cleared at 21,100 *g* at 4 °C for 30 min, and the peptides were desalted with 100 µL Pierce C18 tips (ThermoFisher Scientific, Cat# 87784) using the manufacturer’s protocol. Peptides were then dried in a vacuum centrifuge prior to reconstitution in 0.1% FA. Peptide concentration was estimated with A205 readings on a NanoDrop.

200 ng of peptide digest was separated using a nanoElute nano-flow ultra-high pressure LC system (Bruker Daltonics) coupled online to a trapped ion mobility quadruple-time-of-flight mass spectrometer (timsTOF Pro, Bruker Daltonics) using a CaptiveSpray nano-electrospray ion source. After a 10-min wash at 2% mobile phase B (mobile phase A (MPA): 0.1% FA; mobile phase B (MPB): 0.1% FA in acetonitrile) on a C18 trap column, peptides were separated on a capillary C18 column (25 cm length, 75 µm inner diameter, 1.6 µm particle size, 120 Å pore size; IonOpticks, Fitzroy, VIC, AUS) at a flow rate of 400 nL/min using a stepwise gradient of 2-17% MPB from 10-70 min, 17-25% MPB from 70-100 min, 25-37% MPB from 100-110 min, and 37-85% MPB from 110-120 min, with a 10 min wash at 85% MPB for a total runtime of 130 min. Samples were measured in data-independent analysis - parallel accumulation-serial fragmentation (diaPASEF) mode using 32 windows ranging from 0.6 to 1.421/K_0_ and 400 to 1200 m/z [21].

Raw mass spectrometry (MS) files were processed with DIA-NN in double-pass mode using an *in silico* generated spectral library from the mouse Uniprot reviewed proteome (UP000000589, accessed 20 April 2022) [22]. Peptides were required to be between 200 and 1700 *m/z* with no more than 2 missed cleavages. Cysteine carbamidomethylation was set as a fixed modification, while methionine oxidation and N-terminal acetylation were set as variable modifications. Mass accuracy was set to 15 ppm. Match-between runs was enabled at 1% false-discovery rate (FDR). Heuristic protein grouping was turned off. Protein and peptides identifications were set at a 1% FDR. Protein and peptide output is included in **Table S1**.

Global protein abundance analysis was performed using “DAPAR” [23], “DEP” [24], and “IHW” [25] for R (version 4.1.0) as previously described [26]. Proteins were filtered to remove contaminants and proteins that were not quantified in at least three of five biological replicates within one sample group. Label-free quantification values were Log_2_-transformed and normalized to the median of the total data set. Missing values were imputed via ssla for partially observed values with a sample group or set to 2.5% quantile for values missing across an entire sample group. Limma tests were performed to evaluate statistical significance, *p-values* were adjusted via independent hypothesis weighting based on the number of peptides observed per protein group [25], and statistical significance required a Log_2_-fold change of 0.25 or greater in either direction and an FDR-adjusted *p-value* of 0.05 or less. A list of differentially expressed proteins is included in **Table S2**. Data were further visualized in R. Gene ontology analysis was performed using STRING [27], and the significance threshold was set at an FDR-adjusted *p-value* of 0.05.

### Statistical Analysis

Graphs and statistical analysis were performed with Prism v9.4.0 (GraphPad). Two-way ANOVAs were performed to compare mTORC1 function during development and injury response, followed by Tukey’s multiple comparison tests to detect significant differences between analysis conditions. One-way ANOVAs were performed to compare mTORC1 treatment conditions, followed by multiple comparison tests to detect significant differences between treatment and control conditions. A *p-value* < 0.05 was determined as significant, with all error bars shown as standard error of the mean unless specified.

## RESULTS

### mTORC1 Signaling is Dynamically Regulated During Postnatal Cardiac Development and Following Injury

To define mTORC1 activity in the postnatal heart as well as during cardiac regeneration, we measured the phosphorylation levels of primary downstream targets of mTORC1 during postnatal heart development, as well as during regenerative and non-regenerative models following myocardial infarction (MI) at postnatal day 1 (P1) and P7, respectively. Hearts were collected at 1-day post-surgery (DPS), 7 DPS, and 28 DPS, together with their equivalent timepoints during postnatal development (**Figure 1A**). These timepoints provide a precise timeline of mTORC1 function during postnatal development and following injury by measuring phosphorylation levels of the primary mTORC1 targets, S6K and 4EBP1.

**Figure 1.**
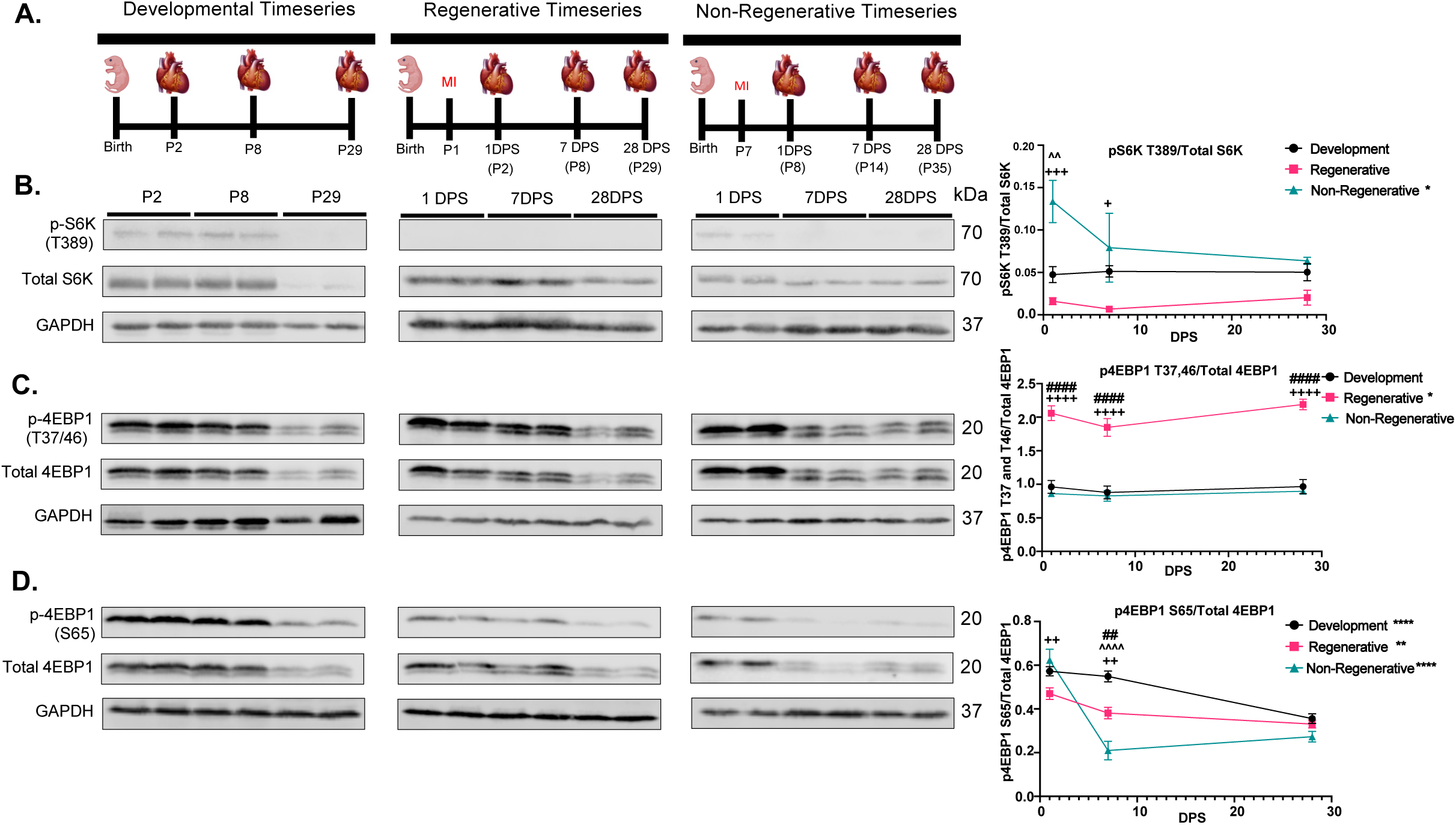
mTORC1 signaling is dynamically regulated during postnatal cardiac development and following injury. **A.** Schematic outlines of developmental collections or post-MI at P1 and P7. **B.** Representative immunoblots and quantification of S6K phosphorylation (pS6K) over total S6K protein during postnatal heart development, regenerative injury response, and non-regenerative injury response, demonstrating that mTORC1 and its downstream target S6K is dynamically regulated during postnatal development and following injury (n=4 per group). **C.** Representative immunoblots and quantification of 4EBP1 (eIF4E Binding Protein 1) rapamycin insensitive phosphorylation sites T37 and T46 over total 4EBP1 protein during development, regenerative injury response, and non-regenerative injury response, demonstrating a significant increase in 4EBP1 T37 and T46 phosphorylation compared to development or non-regenerative hearts (n=4 per group). **D.** Representative immunoblots and quantification of the 4EBP1 rapamycin sensitive phosphorylation site S65 over total 4EBP1 protein, demonstrating dynamic regulation of 4EBP1 S65 phosphorylation depending between development, regenerating, and non-regenerating hearts (n=4 per group). *P<0.05, **P<0.01, ***P<0.001, ****P<0.0001. (*) represents change within treatment between timepoints. (#) represents p-value between development and regenerative conditions. (^) represents p-value between development and non-regenerative conditions. (+) represents p-value between regenerative and non-regenerative conditions.

We quantified a significant decrease in the phosphorylation of S6K during the regenerative injury response, which remains reduced over the course of the timeseries compared to the higher levels of S6K phosphorylation in early postnatal development (**Figure 1B**). This is in stark contrast to non-regenerative injury response, where S6K phosphorylation increases immediately after injury and remains slightly elevated throughout the timeseries. These results demonstrate that mTORC1 phosphorylation of S6K is dynamically regulated during postnatal development and between regenerative and non-regenerative conditions, suggesting that S6K phosphorylation plays an important role following cardiac injury.

mTORC1 phosphorylation of 4EBP1 occurs through sequential phosphorylation of multiple 4EBP1 phosphorylation residues, starting with phosphorylation of the rapamycin insensitive Threonine residues 37 and 46 [28]. We analyzed both rapamycin insensitive and sensitive residues of 4EBP1 by measuring the phosphorylation of 4EBP1-T37/46 and 4EBP1-S65, respectively. We quantified a significant increase in 4EBP1-T37/46 phosphorylation during the regenerative injury response at all timepoints compared to postnatal development and the non-regenerative hearts (**Figure 1C**). Interestingly, phosphorylation of the rapamycin sensitive 4EBP1-S65 residue was distinct between the developmental, regenerative, and non-regenerative hearts (**Figure 1D**). During postnatal development and the regenerative injury response, 4EBP1- S65 phosphorylation increased at 1 DPS and slightly decreased throughout the timeseries, with 4EBP1-S65 phosphorylation levels slightly elevated during postnatal development. However, the non-regenerative hearts exhibit a drastically different response, with 4EBP1-S65 phosphorylation peaking at 1 DPS, then significantly decreasing at 7 DPS, until reaching similar phosphorylation levels as the developmental and regenerative hearts at 28 DPS.

Collectively, our data demonstrate that downstream targets of mTORC1 are dynamically regulated during development, regeneration, and non-regenerative injury response, suggesting that mTORC1 might play an important role during postnatal cardiac regeneration and maturation.

### mTORC1 Inhibition Prevents Neonatal Cardiac Regeneration

To determine whether mTORC1 is required for neonatal cardiomyocyte cell cycle activity and neonatal cardiac regeneration, neonatal mice were treated with the mTOR inhibitors rapamycin (1 mg/kg) or everolimus (1 mg/kg) daily for 3 days at 4, 5, and 6 DPS at P1 (**Figure 2A**). Dosing strategy was determined by optimizing for the dose concentration and treatment duration that resulted in no mortality at baseline (data not shown). Additionally, we utilized this acute treatment protocol as acute rapamycin or everolimus treatments have been demonstrated to be mTORC1 specific with minimal impacts to mTORC2 [29]. To examine whether mTORC1 inhibition alters cardiomyocyte cell cycle activity, we analyzed cardiomyocyte mitosis and cytokinesis at 7 DPS by immunostaining for phospho-Histone H3 and Aurora B Kinase, respectively. We quantified a significant decrease in the number of cardiomyocytes undergoing mitosis and cytokinesis following mTORC1 inhibition (**Figure 2B and C**), suggesting that mTORC1 inhibition blocks cardiomyocyte cell cycle activity, which is the main source of the newly formed cardiomyocytes during neonatal cardiac regeneration.

**Figure 2.**
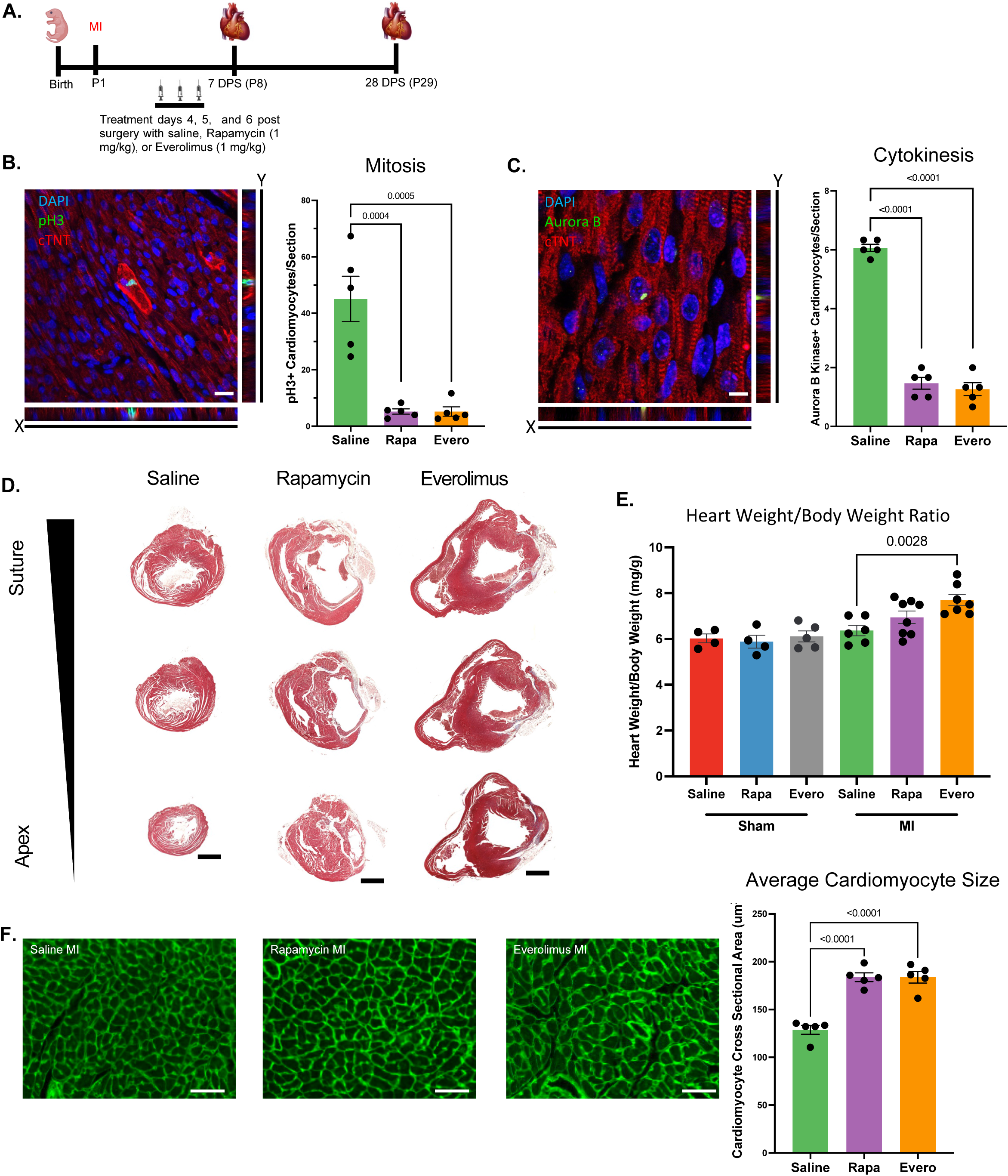
mTORC1 inhibition prevents neonatal cardiac regeneration. **A.** Schematic timeline of MI strategy and rapamycin (Rapa) or everolimus (Evero) treatments. **B.** Immunostaining and quantification of pH3 positive cardiomyocytes demonstrating a significant decrease in cardiomyocyte mitosis following mTORC1 inhibition. **C.** Immunostaining and quantification of Aurora B Kinase positive cardiomyocytes demonstrating a significant decrease in cardiomyocyte cytokinesis following mTORC1 inhibition. **D.** Masson’s Trichrome histological staining of heart section from saline, rapamycin, or everolimus treated mice at 28 days post-MI showing a reduced regenerative capacity after mTORC1 inhibition. **E.** Heart weight to body weight ratio demonstrating a significant increase in heart weight to body weight ratio following mTORC1 inhibition post-MI compared to controls. **F.** Representative images of WGA immunofluorescent staining and cardiomyocyte cross-sectional area quantification showing a significant increase in cardiomyocyte cross-sectional area at 28 days post-MI following mTORC1 inhibition. Quantification of cross-sectional area of 450 cardiomyocytes (150 cardiomyocytes per section for 3 sections) per biological replicate. (Scale bars: B and C, 20 μm; D, 1 mm; F, 50 μm).

To determine whether mTORC1 inhibition of cardiomyocyte proliferation prevents neonatal cardiac regeneration, we performed Masson’s trichrome staining to assess scar formation and myocardial regeneration. Rapamycin and everolimus treated mice exhibited large scar areas along with left ventricular dilation, both hallmarks of loss of cardiac regeneration (**Figure 2D**). We quantified a significant increase in heart weight to body weight ratios in rapamycin and everolimus treated mice post-MI compared to sham controls and saline treated MIs, demonstrating that transient inhibition of mTORC1 by rapamycin or everolimus did not disrupt postnatal development but results in pathological remodeling and hypertrophy post-MI (**Figure 2E, Figure S1A**). Furthermore, we measured cardiomyocyte cross-sectional area by wheat germ agglutinin (WGA) staining. We quantified a significant increase in cardiomyocyte cross sectional area in rapamycin and everolimus treated mice, demonstrating that mTORC1 inhibition promotes cardiac hypertrophy and pathological remodeling post-MI (**Figure 2F**). Interestingly, mTORC1 inhibition did not alter cardiomyocyte cross-sectional area following sham surgery, indicating that the impact of mTORC1 inhibition on cardiomyocyte hypertrophy is an injury-specific response (**Figure S1B**). Collectively, our results demonstrate that mTORC1 function is critical for neonatal cardiomyocyte proliferation and cardiac regeneration.

Since mTORC1 function is necessary for neonatal cardiac regeneration, we wanted to determine the downstream mechanism by which mTORC1 mediates its function. As mTORC1 regulates protein synthesis, we performed the SUnSET assay to measure protein translation in rapamycin and everolimus treated hearts [30, 31]. Following MI at P1, mice were treated with rapamycin or everolimus daily for 3 days at 4, 5, and 6 DPS. This was followed by a single injection with puromycin at 1 hr before harvest at 7 DPS, which can incorporate into newly synthesized proteins. No significant change in puromycin immunoblotting was detected during development or regenerative conditions between saline, rapamycin, or everolimus treated mice, demonstrating that mTORC1 inhibition did not significantly impact protein synthesis (**Figure S2**).

To define the impact of rapamycin and everolimus treatment on mTORC1 function following injury, we analyzed the phosphorylation of S6K and 4EBP1. We quantified a significant reduction in S6K phosphorylation in both rapamycin-and everolimus-treated mice compared to saline-treated controls following sham surgery (**Figure S3A**). As expected, there was no change in phosphorylation of the rapamycin insensitive 4EBP1-T37/46 residue (**Figure S3B**). We also quantified a significant decrease in phosphorylation of the rapamycin sensitive 4EBP1-S65 residue in everolimus-treated mice compared to saline and rapamycin treatments, which may be due to the higher potency of everolimus compared to rapamycin (**Figure S3C**) [32]. Similarly, there was a significant reduction in S6K and 4EBP1-S65 phosphorylation following MI in everolimus-treated mice (**Figure S4A-C**). Taken together, our results demonstrate that mTORC1 is required for heart regeneration, and that this role is not mediated via regulation of protein synthesis but rather through another downstream signaling of mTORC1.

### Global Proteomic Analysis of mTORC1 Inhibition Defines its Role in Cardiac Metabolism

Since transient inhibition of mTORC1 following injury did not disrupt protein synthesis, we sought to investigate the distinct cardiac proteomic signature that drives the loss of regeneration by using a global bottom-up quantitative proteomic analysis. Mice were subjected to sham or MI surgeries at P1, treated intraperitoneally with either saline or rapamycin at 4, 5, and 6 DPS, and then hearts were collected at 7 DPS **(Figure 3A)**. In our analysis, we identified over 6,300 protein groups in each sample group and found that nearly all these protein groups (6,322) were identified in 3 of 5 biological replicates in all four sample groups **(Figure 3B)**, which confirms our earlier data that rapamycin treatment does not drastically alter gross proteome composition. Furthermore, the proteomic data showed a high degree of normality and median alignment **(Figure S5A)**, low coefficients of variation among sample groups **(Figure S5B)**, and high data completeness **(Figure S5C)**, bolstering our confidence in downstream quantitative analyses. When examining our samples using a dimensional reduction analysis, we noticed the largest separation among sample groups was due to injury (separation along PC1) **(Figure 3C)**. It is clear, however, that rapamycin treatment also affects proteome composition, as saline and rapamycin-treated samples cluster separately **(Figure 3C)**. Therefore, we first examined how mTORC1 inhibition affects baseline cardiac proteome composition.

**Figure 3.**
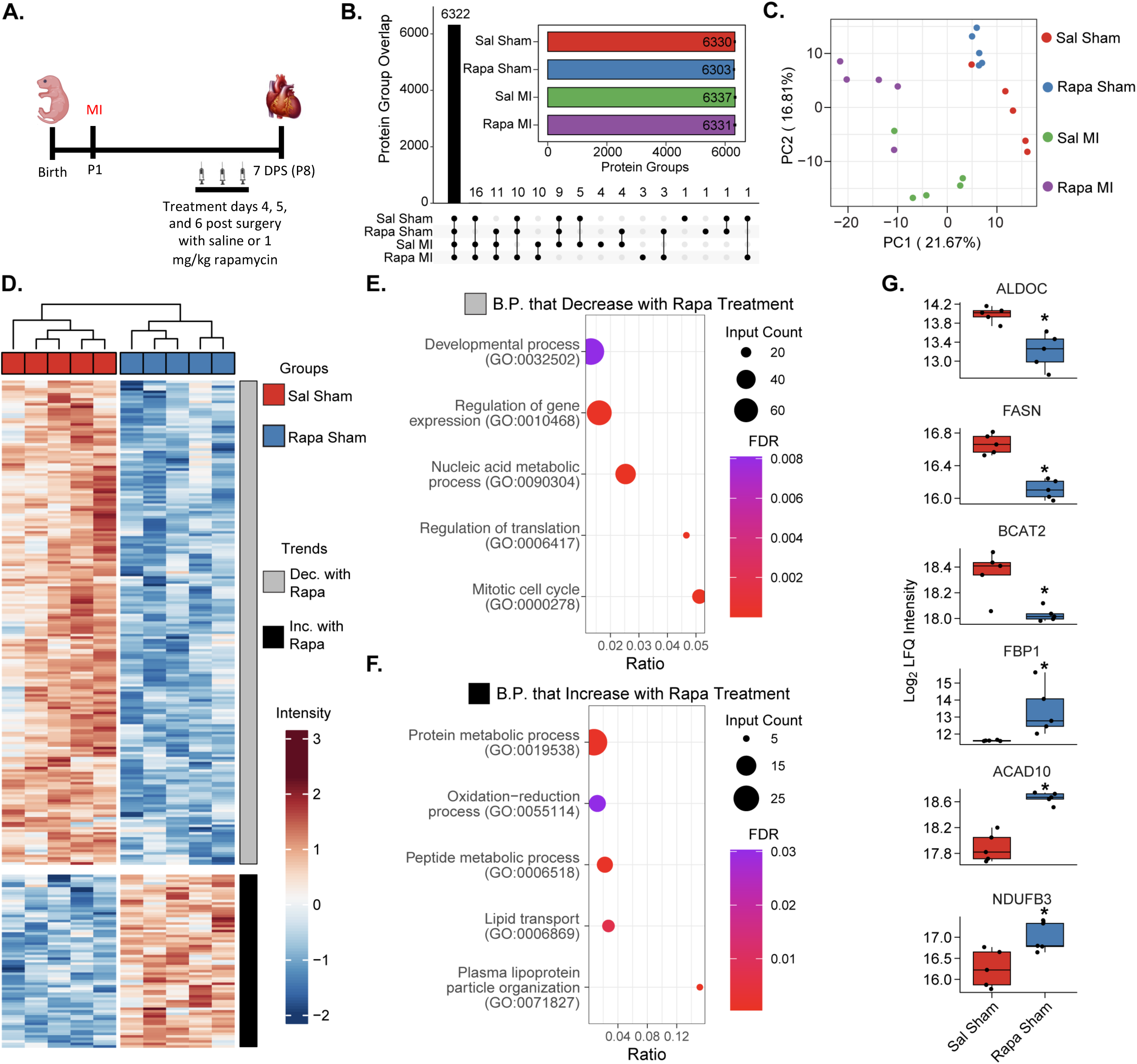
Global proteomic analysis of mTORC1 inhibition highlights its role in cardiac metabolism. **A.** Outline of experimental design. Mice at P1 (n=5 per group) were given sham or myocardial infarction (MI) surgeries. Mice were treated 4, 5, 6 days post-surgery (DPS) with I.P. injection of saline (Sal) or rapamycin (Rapa). Hearts were harvested for global proteomic analysis at 7 DPS. **B.** Protein IDs were filtered to require n ≥ 2 per sample group for representation. Inlay: Bar chart demonstrating unique protein groups identified from each sample group (bar demonstrates mean, error bars demonstrate s.e.m.). **C.** Principal component analysis of per-sample Log_2_ protein abundances demonstrates separation of sample groups based on treatment (Sal or Rapa) and surgery (sham or MI). **D.** Hierarchal heatmap demonstrating *z-score* normalized intensities of significantly differently expressed proteins between saline and rapamycin treated sham mice (adjusted p-value ≤ 0.05 and |Log_2_ Fold Change| ≥ 0.25 required to be considered significant). The heatmap is separated into two clusters: proteins that decrease in expression with rapamycin treatment (grey) and proteins that increase with rapamycin treatment (black). **E. and F.** STRING Biological process gene ontology (GO) analysis of proteins that decrease in expression with rapamycin treatment (**E**) and proteins that increase in expression with rapamycin treatment (**F**). Dot size represents the number of identified proteins within a GO group. Gene ratio indicates the fraction of all proteins within the GO group that were identified. Color represents the FDR-adjusted p-value of the GO overrepresentation test. **G.** Boxplots displaying key proteins significantly dysregulated by rapamycin treatment in sham mice. Boxplots represent n=5 biological replicates per experimental group (* indicates adjusted p-value ≤ 0.05 and |Log_2_ Fold Change| ≥ 0.25).

Using a limma-based differential expression analysis, we were able to identify 262 differentially expressed proteins (DEPs) between saline and rapamycin-treated sham P8 mouse hearts **(Table S2)**. When plotting these DEPs on a *z-score* normalized heatmap, we noticed that sample groups clustered together and there were two distinct trends: a group of proteins that decreased with rapamycin treatment and a group of proteins that increased **(Figure 3D)**. Interestingly, rapamycin treatment led to a larger cluster of proteins with both decreased and increased abundance when compared to saline-treated mice **(Figure 3D)**. Using the group of downregulated proteins, we performed gene ontology (GO) analysis for biological processes with STRING and observed that rapamycin-treatment led to a decrease in abundance of proteins involved in cell cycle and translational regulation processes **(Figure 3E, Table S3)**, including regulation of gene expression (GO: 0010468), nucleic acid metabolic processes (GO: 0090304), regulation of translation (GO: 0006417), and mitotic cell cycle (GO: 0000278). On the other hand, GO analysis of the upregulated proteins following rapamycin-treatment demonstrate an increase in level of proteins involved in metabolism including oxidative-reduction processes (GO:0055114) and lipid transport (GO:0006869) (**Figure 3F, Table S3**). This indicates that under baseline conditions, systemic administration of rapamycin leads to changes in protein levels related to cell cycle and metabolism.

Specifically, we noted a change in protein expression that indicates an increase in cardiomyocyte metabolic maturation after mTORC1 inhibition compared to control hearts. Of note, there was a significant reduction in the protein levels of glycolysis and branched chain amino acid (BCAA) catabolism enzymes such as ALDOC, FASN, and BCAT2 following mTORC1 inhibition (**Figure 3G**). Additionally, there was a significant increase in multiple mitochondrial metabolism proteins including ACAD10 and NDUFB3 **(Figure 3G)**. Our proteomic analysis indicates that mTORC1 inhibition by rapamycin in the postnatal heart shifts the neonatal proteomic landscape from glycolysis and BCAA catabolism towards increased oxidative phosphorylation that may result in premature metabolic maturation. These results suggest that mTORC1 inhibition via rapamycin treatment shifts the cardiac proteomic landscape towards a more mature metabolic state without inducing hypertrophic remodeling in the uninjured heart (**Figure 2E, Figure S1**).

### mTORC1 Inhibition Prevents Cardiac Regeneration Through Metabolic Remodeling

Since we identified distinct separation between saline and rapamycin-treated mice post-MI at 7 DPS in our principal component analysis **(Figure 3C)**, we next analyzed how rapamycin-treatment affects proteome composition after injury to glean insight into how mTORC1 inhibition prevents cardiac regeneration **(Figure 3A)**. We identified 251 DEPs, with 117 proteins upregulated in the saline-treated regenerative mice and 134 proteins upregulated in the rapamycin-treated non-regenerative mice **(Figure 4A)**. Using the proteins that were upregulated in the saline-treated regenerative mice, we identified GO processes related to nucleotide biosynthetic processes (GO: 0009165), positive regulation of cardiac muscle tissue growth (GO: 0055023), and mitotic cell cycle (GO: 0000278) **(Figure 4B)**. As cardiac regeneration is a function of cardiac development [33], it is plausible that these developmental processes would be upregulated during a regenerative response to injury. Conversely, when we perform GO analysis on the proteins upregulated in the rapamycin-treated non-regenerative mice, we identified processes related to regulation of cell death (GO: 0010941) and Fibrinolysis (GO: 0042730) **(Figure 4C, Table S4)**, indicating a pathological response to injury. Interestingly, we also detected an increase in expression of proteins associated with oxidative phosphorylation in the rapamycin-treated non-regenerative mice, including oxidative-reduction processes (GO: 0055114) and fatty acid metabolic processes (GO: 0006631) **(Figure 4C, Table S4)**, further underlining the role of mTORC1 in cardiac metabolism and its influence over cardiac regeneration.

**Figure 4.**
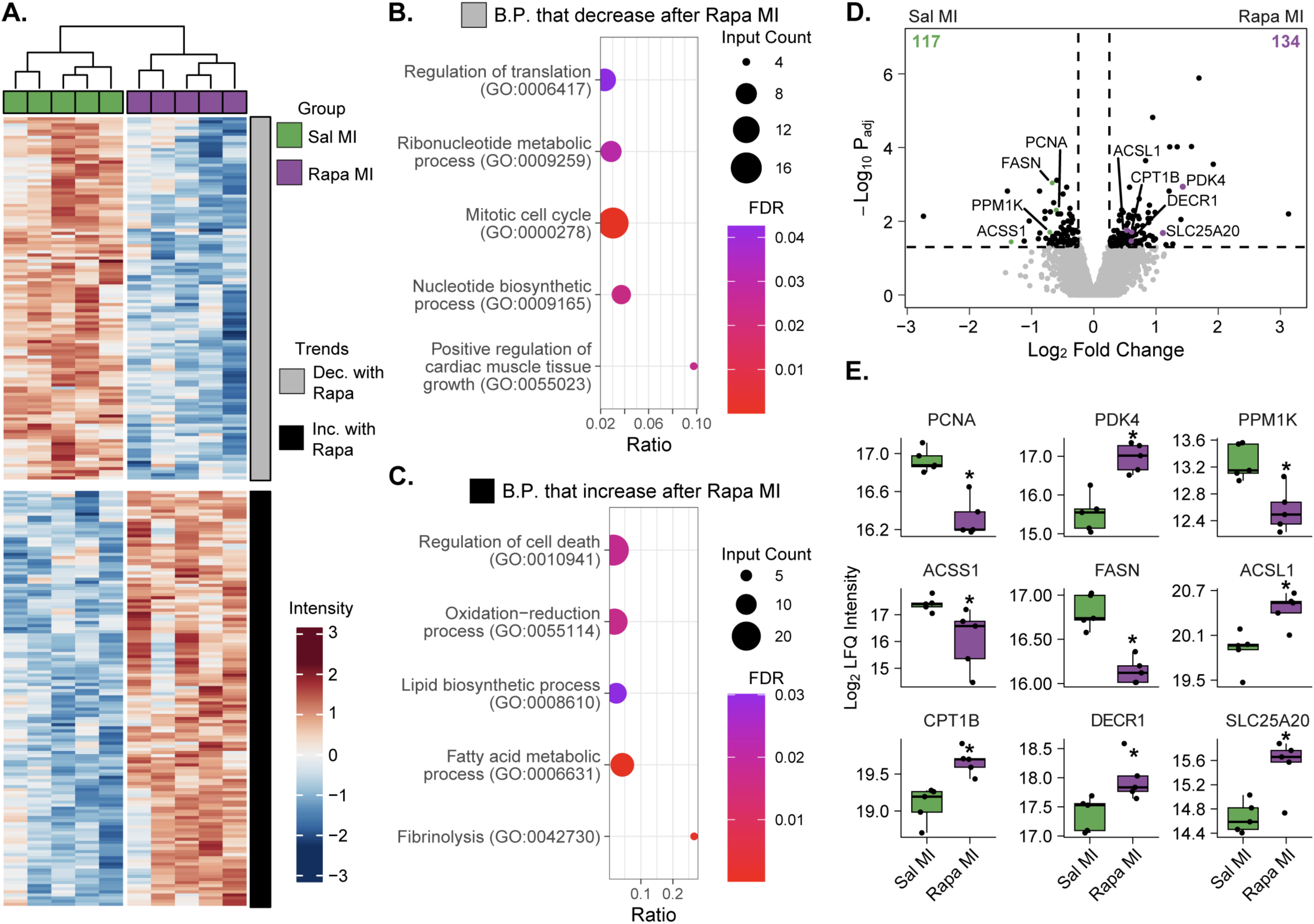
mTORC1 inhibition prevents cardiac regeneration through a metabolic remodeling. **A.** Hierarchal heatmap demonstrating *z-score* normalized intensities of significantly differently expressed proteins between saline and rapamycin-treated MI mice (adjusted p-value ≤ 0.05 and |Log_2_ Fold Change| ≥ 0.25 required to be considered significant). The heatmap is separated into two clusters: proteins that decrease in expression with rapamycin treatment (grey) and proteins that increase with rapamycin treatment (black). **B and C.** STRING Biological process gene ontology (GO) analysis of proteins that decrease in expression with rapamycin treatment after MI (**B**) and proteins that increase in expression with rapamycin treatment post-MI (**C**). Dot size represents the number of identified proteins within a GO group. Gene ratio indicates the fraction of all proteins within the GO group that were identified. Color represents the FDR-adjusted p-value of the GO overrepresentation test. **D.** Volcano plot displaying fold-change in protein expression between saline and rapamycin-treated MI mice. The number of significantly upregulated proteins per group is shown in the bottom corners of the comparison (n = 5 per group, * indicates adjusted p-value ≤ 0.05 and |Log_2_ Fold Change| ≥ 0.25). **E.** Boxplots displaying key proteins significantly dysregulated by rapamycin treatment after MI. Boxplots represent n=5 biological replicates per experimental group (* indicates adjusted p-value ≤ 0.05 and |Log_2_ Fold Change| ≥ 0.25).

Of specific interest, we identified differential expression of key cardiac and metabolic proteins between the non-regenerating rapamycin-treated mice and the saline-treated regenerative controls **(Figure 4D and 4E)**. Regeneration is dependent on a glycolytic metabolic state, which is supported by a significant decrease in PDK4 levels, demonstrating an increase in glycolytic metabolism in saline-treated mice compared to rapamycin-treated mice. Rapamycin-treated hearts displayed reduced levels of PPM1K, ACSS1, and FASN compared to saline-treated controls, indicating a reduced rate of BCAA metabolism and lipid synthesis in the non-regenerative hearts. Furthermore, rapamycin treatment results in a significant upregulation of oxidative phosphorylation and fatty acid oxidation proteins post-MI, such as ACSL1, CPT1B, DECR1, and SLC25A20. This increase in oxidative phosphorylation proteins in the rapamycin-treated hearts is concomitant with a significant decrease in protein levels of the proliferation marker PCNA, supporting our results that rapamycin-treatment reduces cardiomyocyte proliferation and heart regeneration following MI (**Figure 2**). These results demonstrate that mTORC1 inhibition by rapamycin reduces the protein levels of glycolytic enzymes while increasing the protein levels of oxidative phosphorylation and fatty acid oxidation enzymes following injury, suggesting that mTORC1 inhibition accelerates the metabolic maturation of cardiomyocytes that may contribute to the loss of cardiomyocyte proliferation and heart regeneration post-MI.

## DISCUSSION

The postnatal metabolic switch of the mammalian heart from glycolysis to fatty acid oxidation is an essential part of cardiac maturation and maintenance of adult heart function; however, this switch contributes to loss of the endogenous regenerative potential of the mammalian heart. Recent studies demonstrate that inhibition of fatty acid oxidation and mitochondrial oxidative phosphorylation via pyruvate dehydrogenase kinase 4 (PDK4) deletion or inhibition of succinate dehydrogenase (SDH) can induce adult cardiomyocyte proliferation and heart regeneration [34, 35]. Thus, elucidating the mechanisms that control this metabolic switch can lead to the development of novel targeted approaches to promote cardiac regeneration.

mTORC1 represents a central node in regulating metabolic pathways and protein synthesis that influence cellular growth as well as cell fate [8]. However, the role of mTORC1 during postnatal cardiac maturation and its influence on heart regeneration is unclear. In this study, we demonstrate that mTORC1 signaling is dynamically regulated during postnatal development and regeneration in contrast to the non-regenerating heart. These results suggest that mTORC1 may play an important role during cardiomyocyte proliferation and regeneration. Interestingly, mTORC1 inhibition by treatment with either rapamycin or the rapalog, everolimus, impedes cardiomyocyte proliferation and blocks neonatal mouse heart regeneration. Although the overall levels of protein synthesis were not affected by transient inhibition of mTORC1, our proteomic analysis demonstrates that acute mTORC1 inhibition by rapamycin treatment results in a distinct proteomic signature, which exhibits reduced levels of key glycolytic enzymes and higher levels of oxidative phosphorylation and fatty acid oxidation proteins suggesting that mTORC1 inhibition promotes postnatal metabolic maturation. This is evident by the reduced levels of myocyte proliferation and inhibition of heart regeneration following injury. These results suggest that mTORC1 inhibition specifically accelerates metabolic maturation post-MI compared to sham controls, which reduces rates of cardiomyocyte proliferation and increases cardiomyocyte size following infarction. This is consistent with a recent study that demonstrates that mTORC1 inhibition can promote maturation of induced pluripotent stem cell derived cardiomyocytes *in vitro* [36].

Our results suggest that targeting mTORC1 could be a promising strategy for modulating the cardiomyocyte transition from a hyperplastic to a hypertrophic state to promote heart regeneration following injury. Furthermore, mTOR signaling may represent an evolutionarily conserved mechanism during regeneration, as a recent study demonstrates an important role for mTOR activity during zebrafish heart regeneration [37]. However, the downstream metabolic state following mTORC1 inhibition remains to be fully defined. Elucidating how mTORC1 impacts cardiomyocyte metabolism specifically with respect to oxidative phosphorylation and fatty acid synthesis is important to fully establish the role of mTORC1 on cardiomyocyte maturation and function. Furthermore, understanding the role of mTORC1 in multiple cell types during heart regeneration needs to be determined. In addition, rapamycin and everolimus are known immunosuppressants, thus whether this inhibition of myocyte proliferation and regeneration is partially mediated via targeting the immune response needs to be established. Nevertheless, our current study highlights the importance of mTORC1 in regulating mammalian cardiomyocyte cell cycle activity and heart regeneration, suggesting that mTORC1 could be a promising therapeutic target for heart failure.

## Author contributions

W.G.P., T.J.A., J.B. performed the experiments and prepared the manuscript. W.G.P., T.J.A., K.A.H., A.I.M. designed the experiments, performed the analyses, and wrote the manuscript. D.J.N., C.P., K.W., R.N., contributed to western blotting, imaging, and quantification. T.J.A., Y.G. performed the global proteomics and analyses and edited the manuscript. A.I.M. conceived the project and wrote the manuscript.

## Data availability

Source data are available via the MassIVE repository with identifier MSV000092436 and the PRIDE repository via ProteomeXchange with identifier PXD043771. All data are available from the corresponding author upon request.

## Supporting information

Supplemental Datasets for Western

## Acknowledgements

Funding for this project was provided by NIH/NHLBI under Ruth L. Kirschstein NRSA T32 HL007936 to the UW Cardiovascular Research Center (W.G.P.), NIH/NIGMS Training Program in Molecular and Cellular Pharmacology T32 GM008688 (T.J.A.), AHA Career Development Award 19CDA34660169 (A.I.M.), NIH/NHLBI R56 HL155617 (A.I.M.), NIH/NHLBI R01 HL166256 (A.I.M.), DOD W81XWH2210094 (A.I.M.). Y.G. would like to acknowledge NIH/NHLBI R01 HL109810, R01 HL096971, and S10 OD018475. We thank Lance Rodenkirch and the Optical Imaging Core grant S10 OD025040-01 for imaging support. We thank members of the Mahmoud lab for critical reading of the manuscript.

