## Supplementary material for "mTORC1 Regulates the Metabolic Switch of Postnatal Cardiomyocytes During Regeneration": Figures S1-S5

**Figure S1**

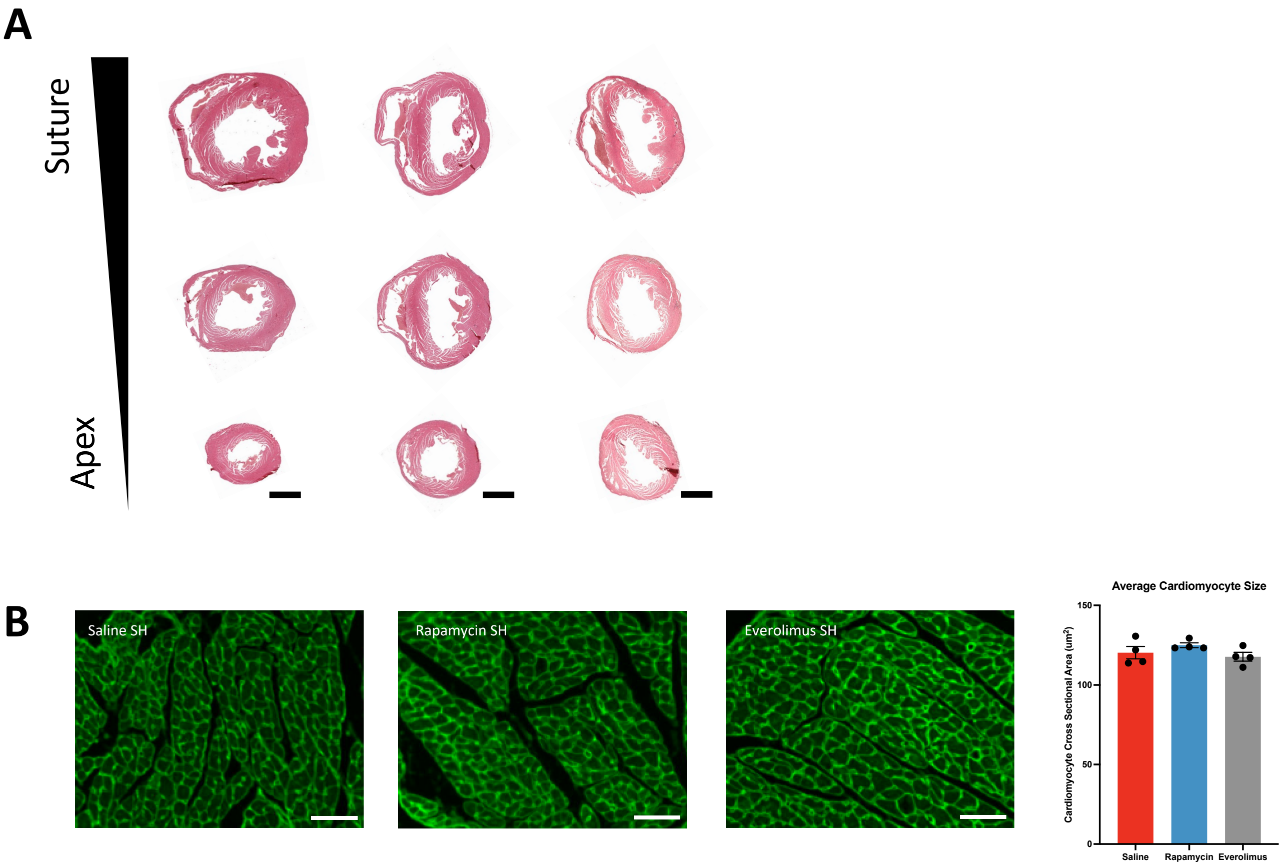


**Figure S1. Histological analysis following rapamycin and everolimus treatment post sham surgery. A.** Masson's trichrome histological staining of heart sections at P29 from saline, rapamycin, or everolimus treated mice. **B.** Representative images of wheat germ agglutinin (WGA) immunofluorescent staining and quantification of cardiomyocyte cross sectional area showing no significant difference in cardiomyocyte size between groups. Quantitative analysis of 150 cardiomyocytes across 3 sections for 450 total cardiomyocytes measured per biological replicate. (Scale bars: A, 1 mm; B, 50 um).

**Figure S2**

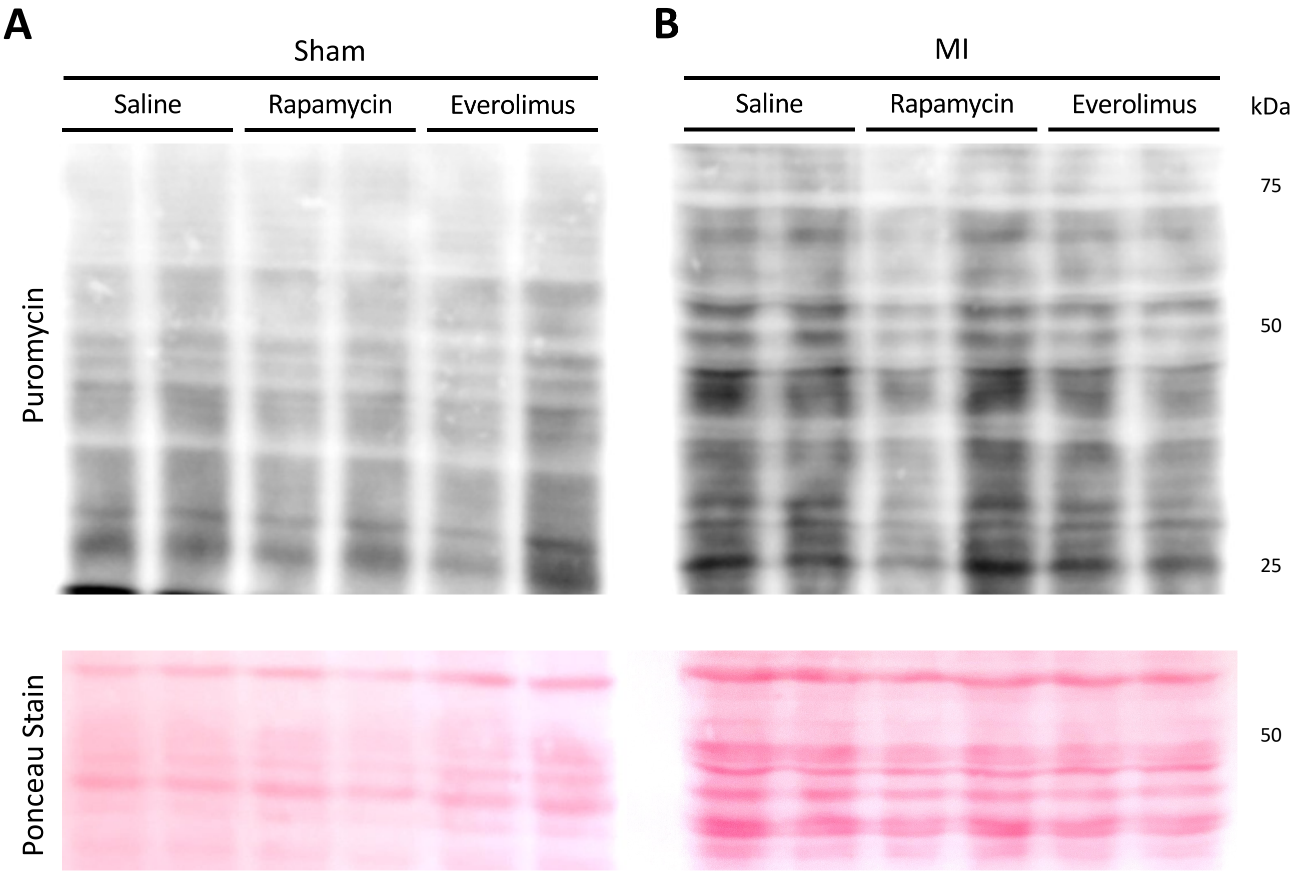


**Figure S2. SUnSET analysis of translational rate following rapamycin or everolimus treatment. A**. Upper, Puromycin stain showing total protein translation at 1 hour before heart collection at 7 dps SH. Lower, Ponceau stain acting as loading control. **B**. Upper, Puromycin stain showing total translation at 1 hour before heart collection at 7 dps MI. Lower, Ponceau stain acting as loading control.

### Figure S3

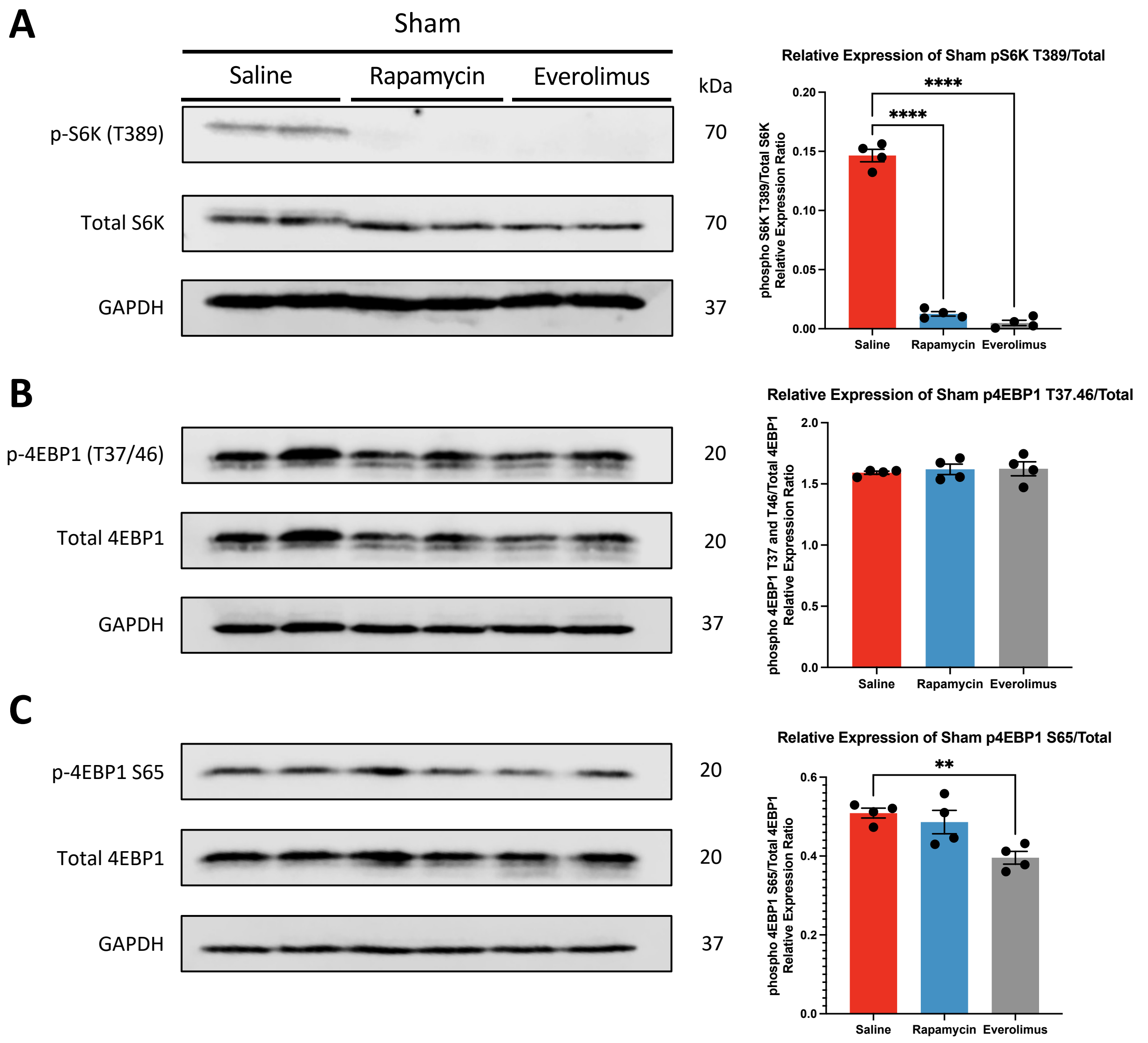


**Figure S3. mTORC1 signaling at 7 days post sham surgery following rapamycin or everolimus treatment. A.** Representative immunoblots and quantification of S6K activation under sham surgical conditions following treatment with saline, rapamycin, or everolimus, showing significant decrease of S6K T389 phosphorylation after mTORC1 inhibition. **B.** Representative immunoblots and quantification of 4EBP1 (eIF4E Binding Protein 1) rapamycin insensitive phosphorylation sites T37 and T46 showing no significant differences between all conditions. **C.** Representative immunoblots and quantification of 4EBP1 rapamycin sensitive phosphorylation site S65 showing a significant decrease in 4EBP1 S65 phosphorylation after everolimus treatment. **P<0.01, ****P<0.0001.

**Figure S4**

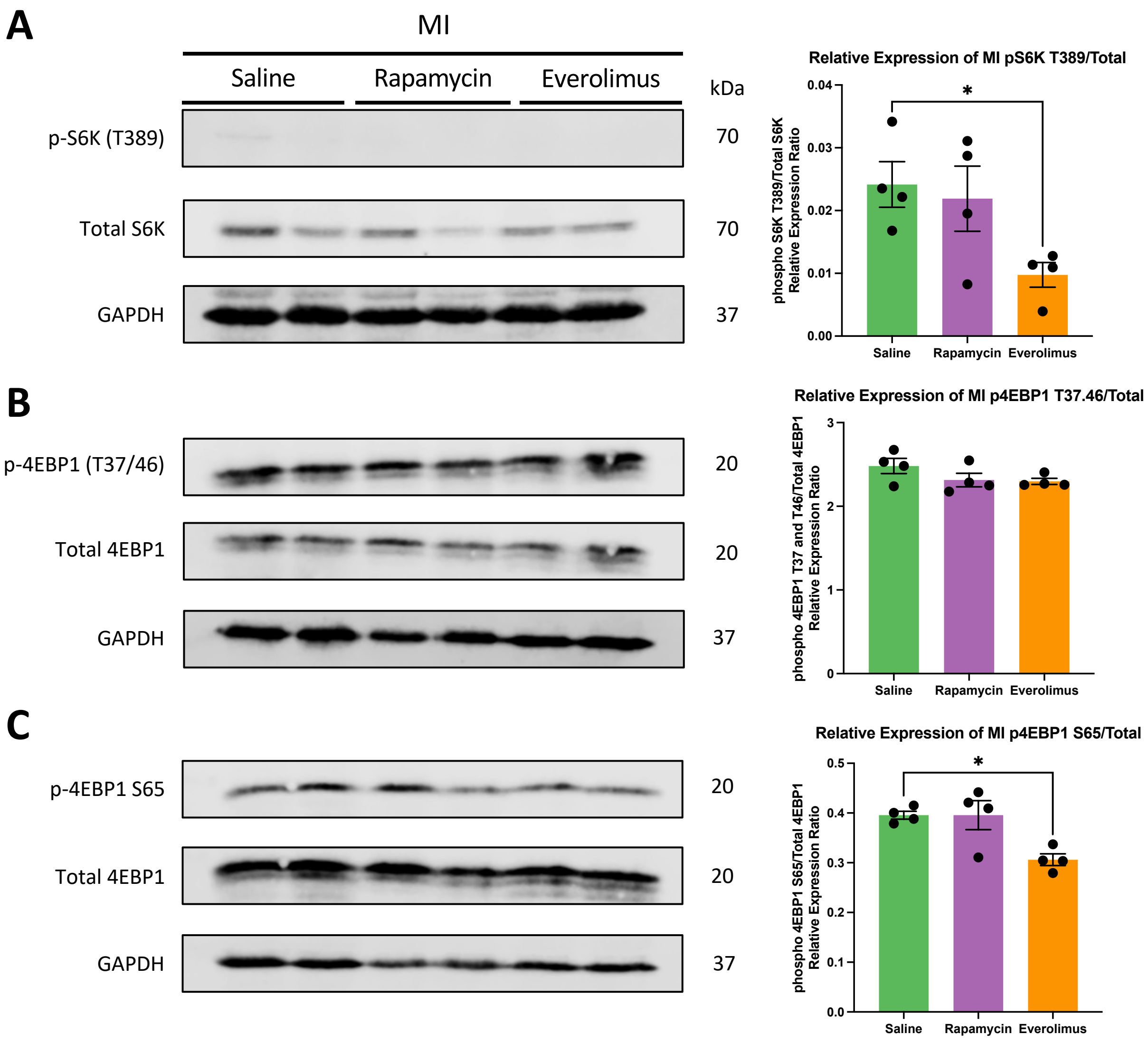


**Figure S4. mTORC1 signaling at 7 days post myocardial infarction following rapamycin or everolimus treatment. A.** Representative immunoblots and quantification of S6K activation post-MI following saline, rapamycin, or everolimus treatment, showing slight reduction of S6K T389 phosphorylation after mTORC1 inhibition. **B.** Representative immunoblots and quantification of 4EBP1 rapamycin insensitive phosphorylation sites T37 and T46 showing no significant changes between conditions. **C.** Representative immunoblots and quantification of 4EBP1 rapamycin sensitive phosphorylation site S65 showing a significant decrease in 4EBP1 S65 phosphorylation after everolimus treatment. *$P<0.05$.

**Figure S5**

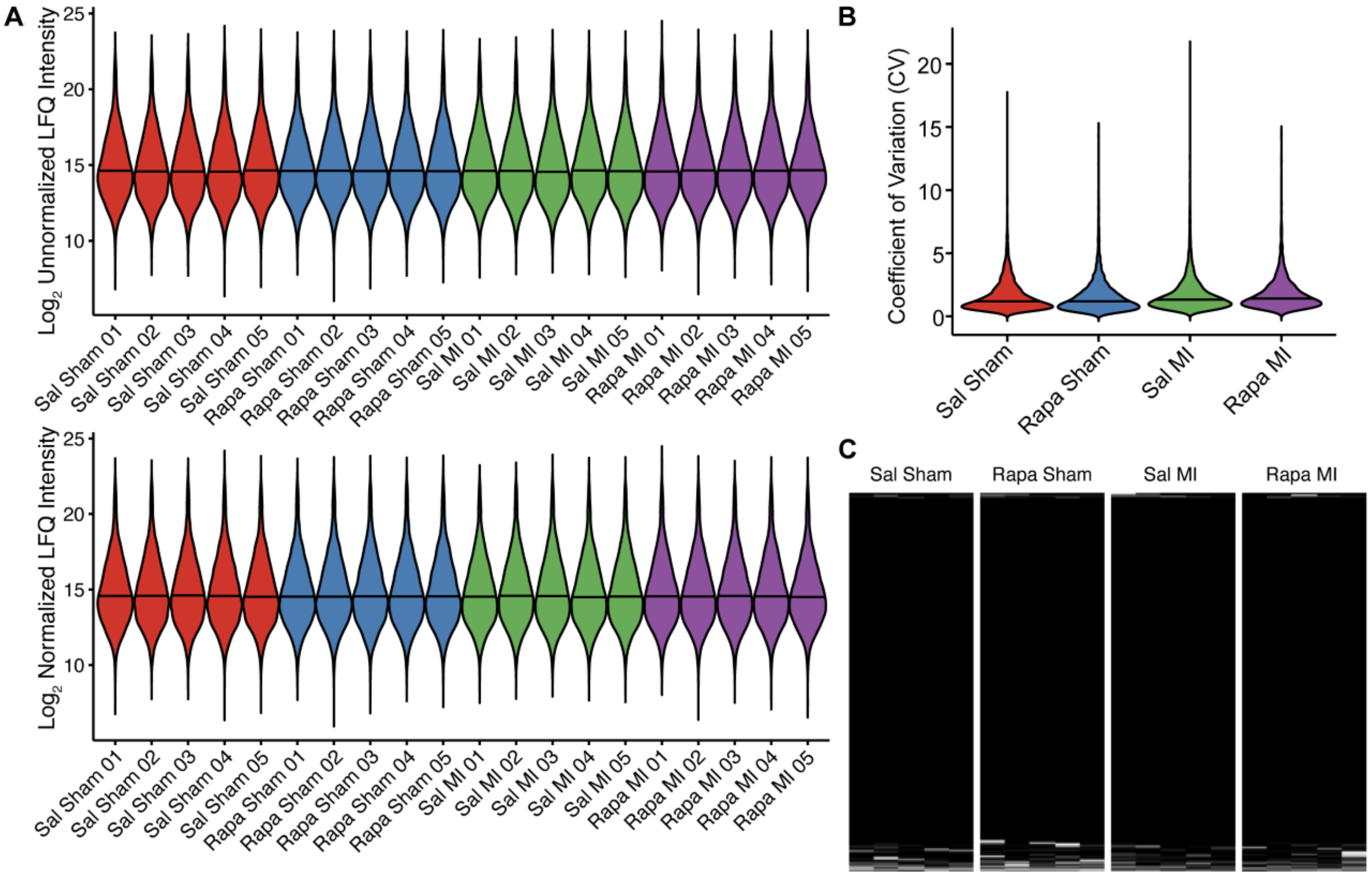


**Figure S5. Bottom-up proteomics quality control metrics. A.** Violin plots demonstrating $Log_2$ unnormalized and median normalized LFQ protein group intensity. The line indicates the sample median. The striking similarity between the unnormalized and normalized data highlights the reproducibility of sample preparation and data acquisition. **B.** Violin plots showing the coefficient of variation among protein groups within each biological group (n = 5 per group). The line indicates the sample median. **C**. Missing value plot demonstrating high data completeness before imputation. White indicates a missing value.
