## Supplemental Datasets for Western for "mTORC1 Regulates the Metabolic Switch of Postnatal Cardiomyocytes During Regeneration"

### Figure 1B: Development

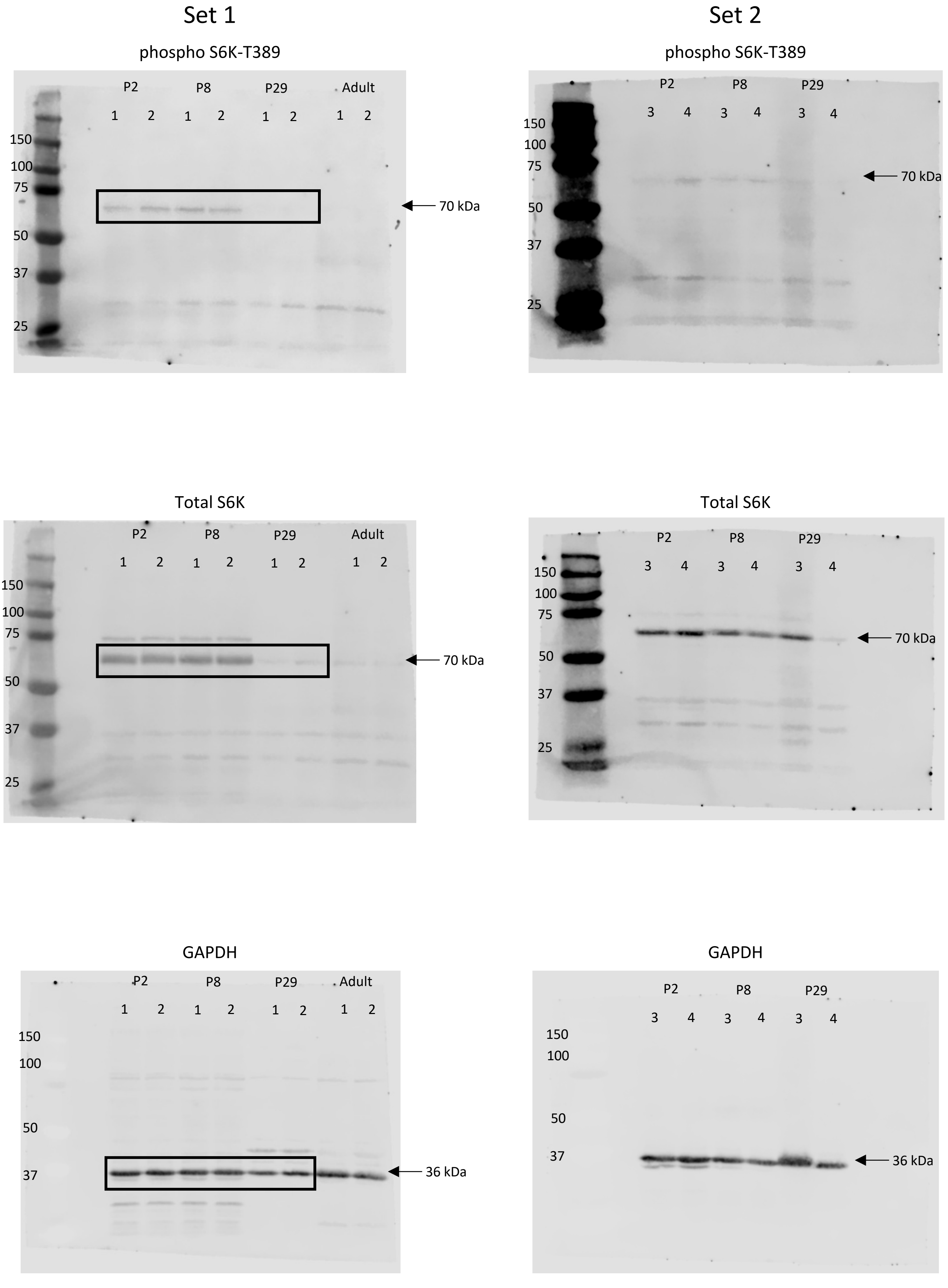

### Figure 1B: Regenerative

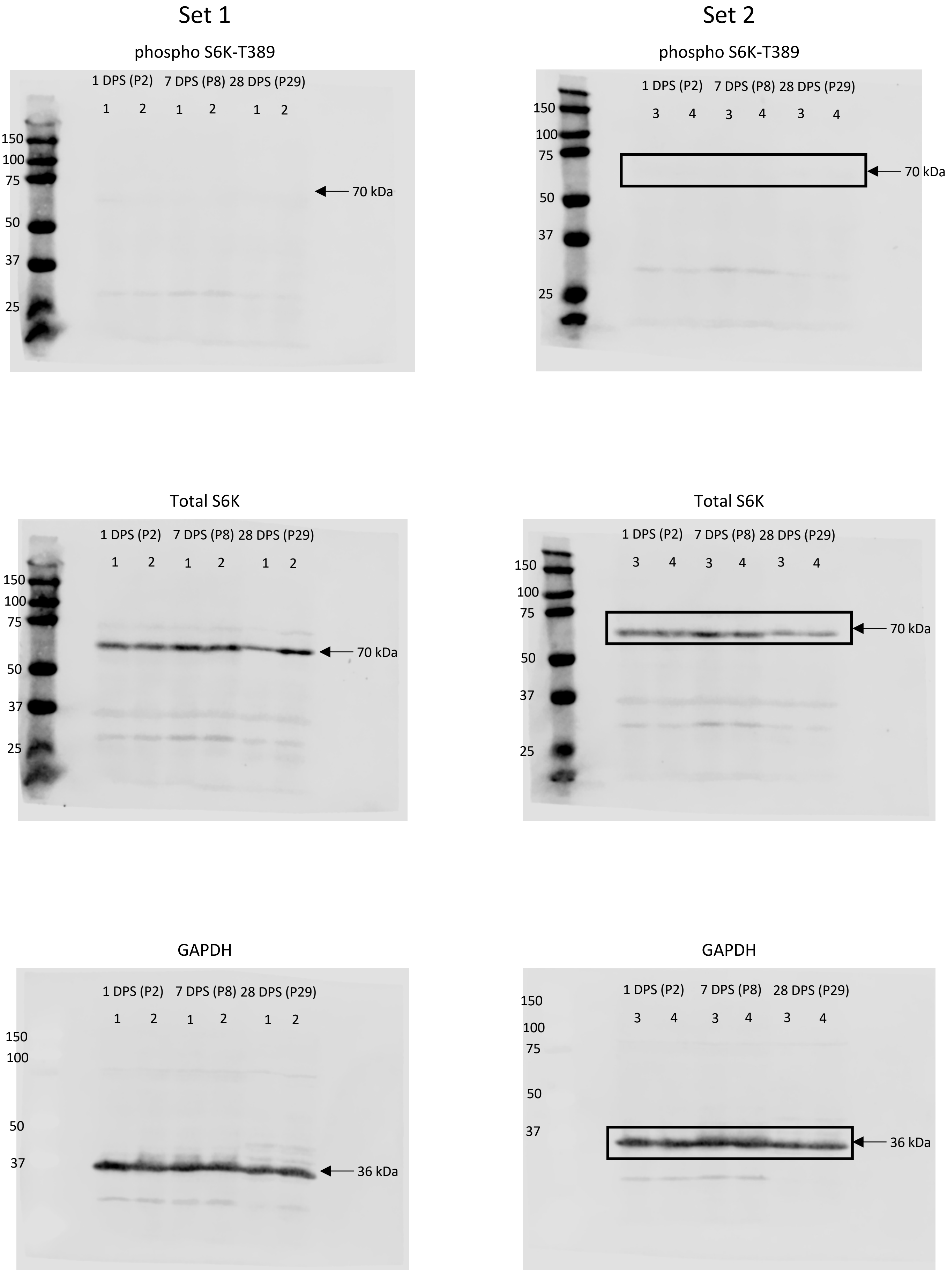

### Figure 1B: Non-Regenerative

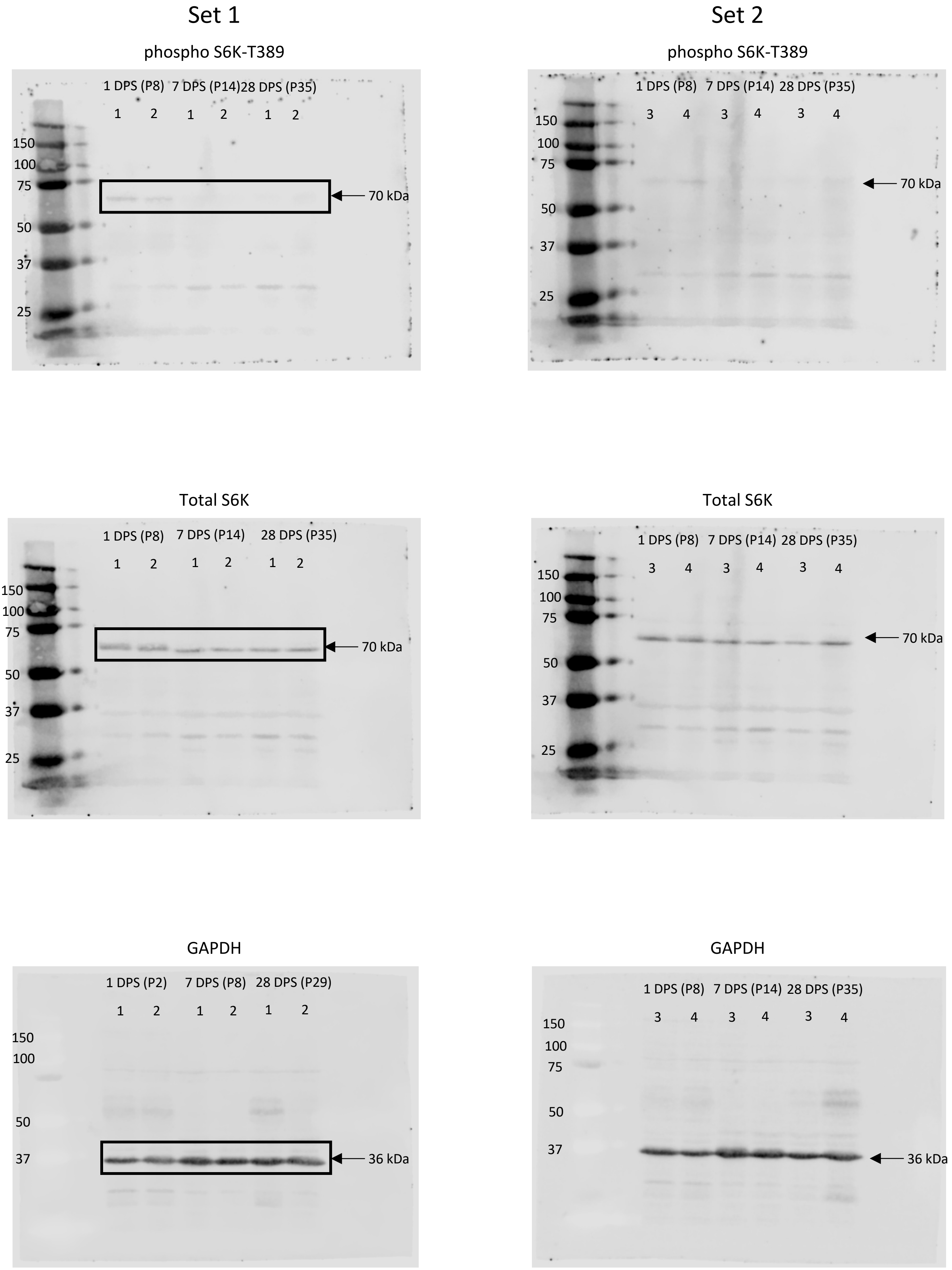

### Figure 1C: Development

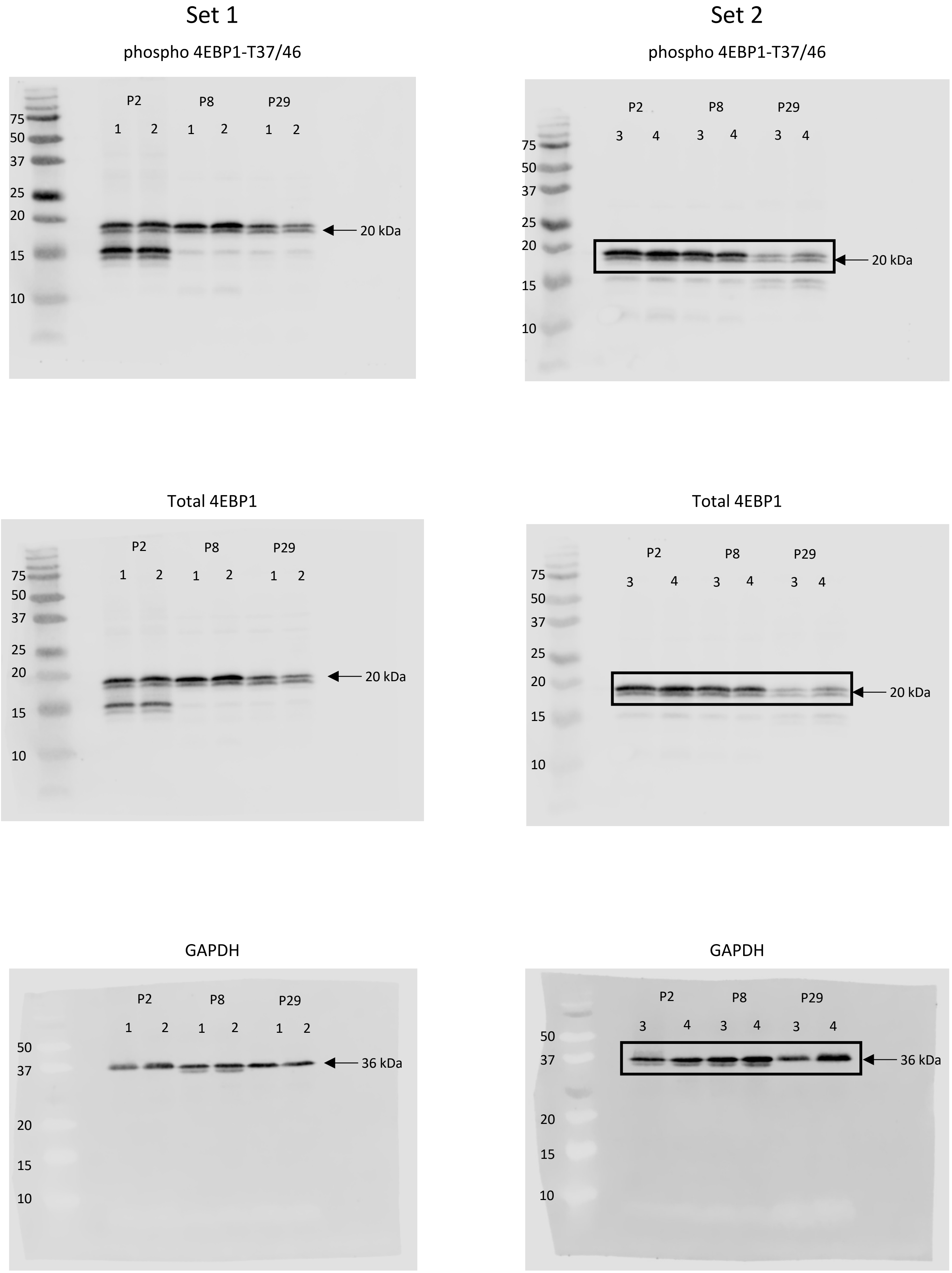

### Figure 1C: Regenerative

Set 1

phospho 4EBP1-T37/46

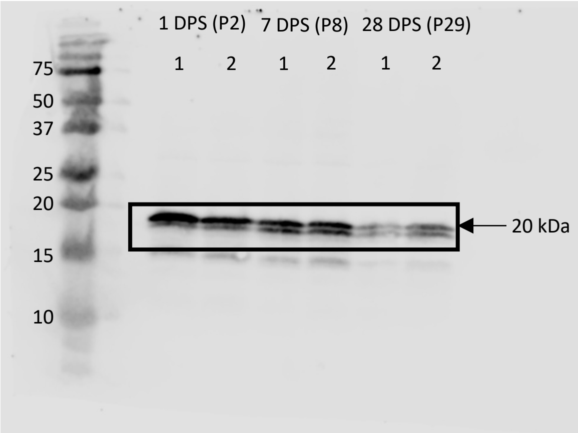


Set 2

phospho 4EBP1-T37/46

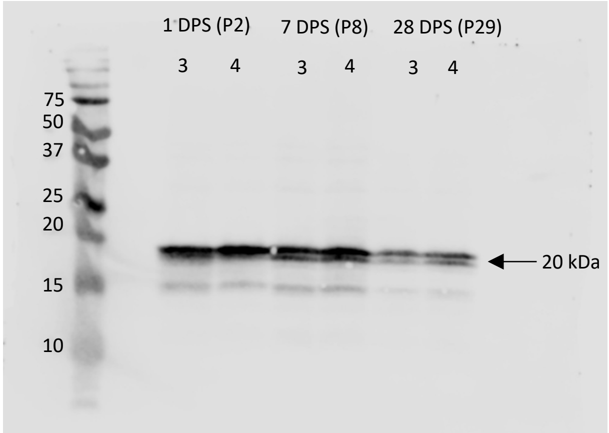


Total 4EBP1

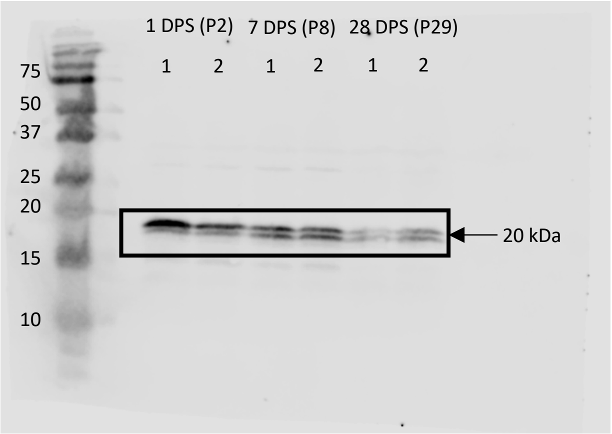


Total 4EBP1

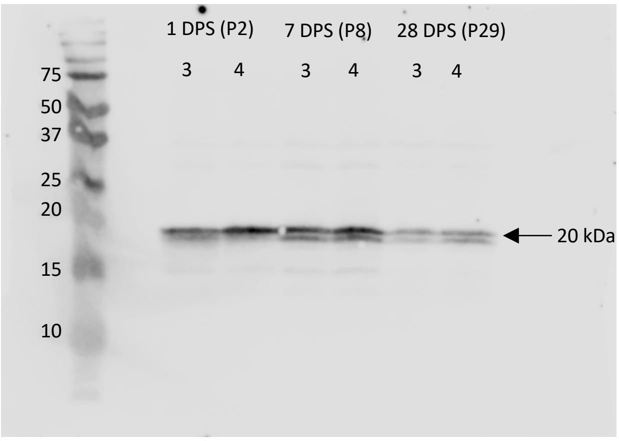


GAPDH

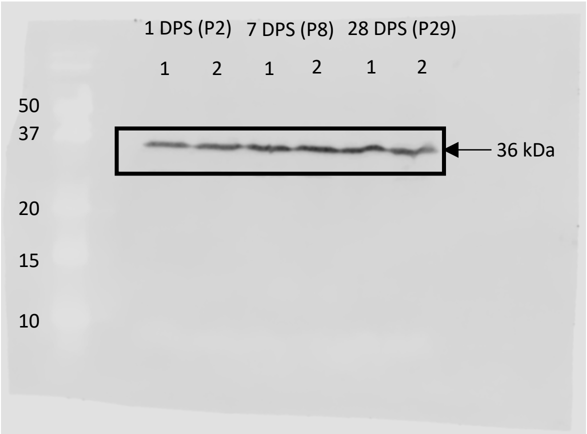


GAPDH

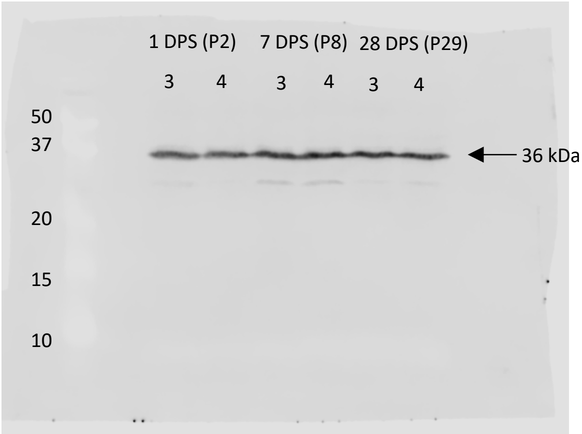

### Figure 1C: Non-Regenerative

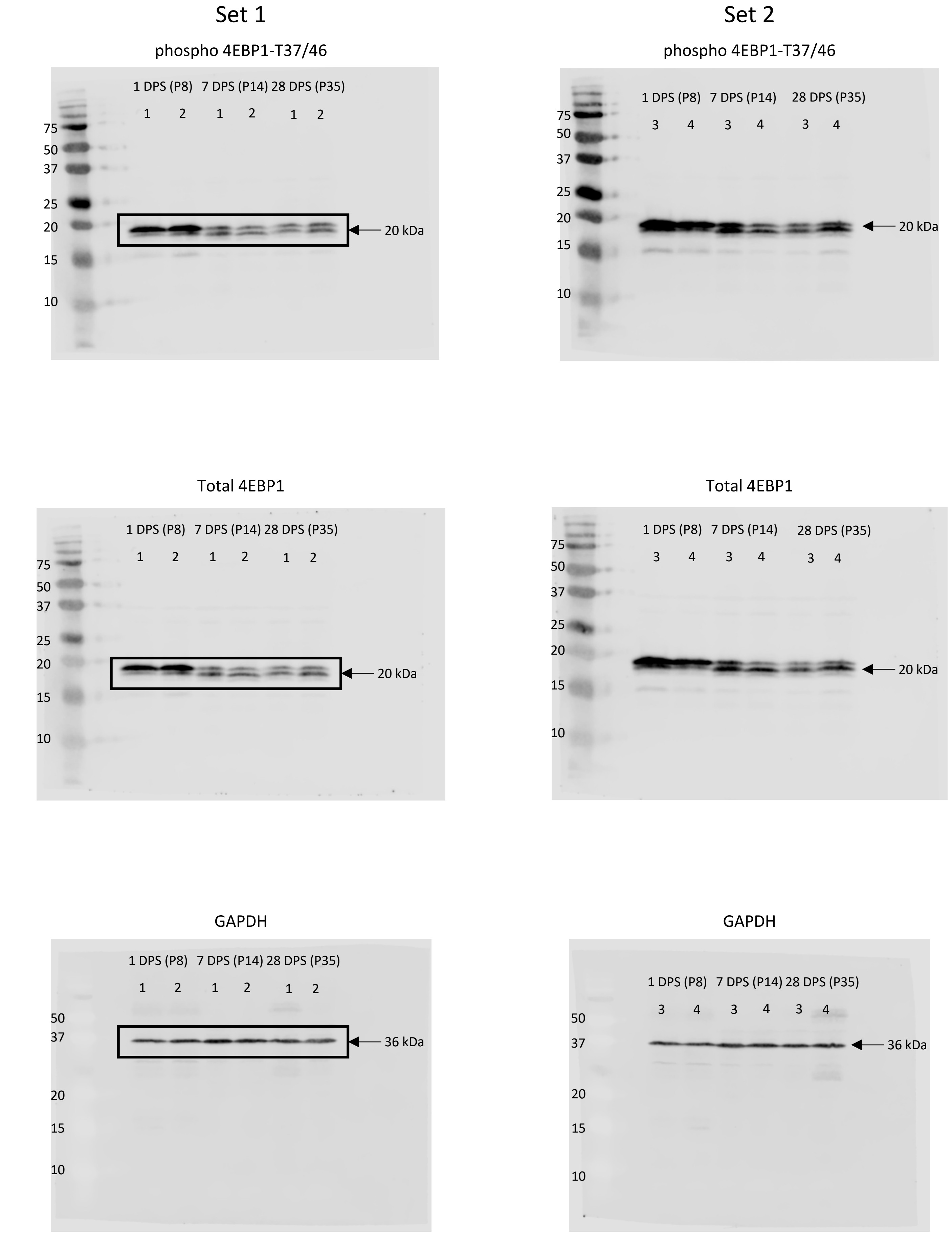

### Figure 1D: Development

#### Set 1

phospho 4EBP1-S65

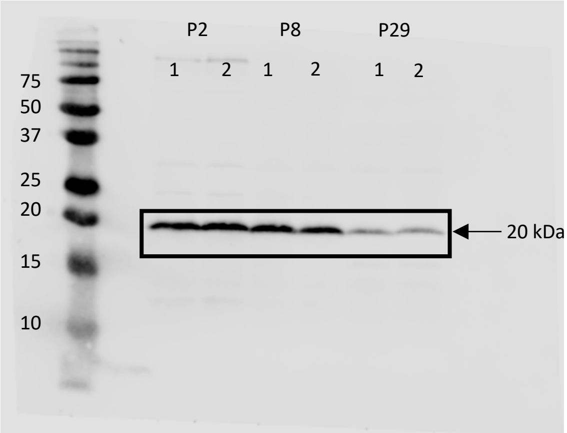


Total 4EBP1

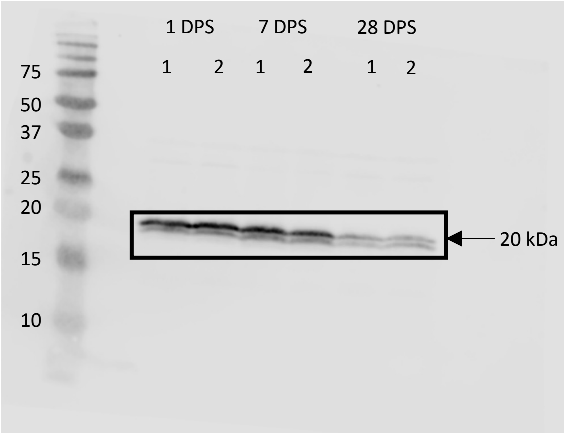


GAPDH

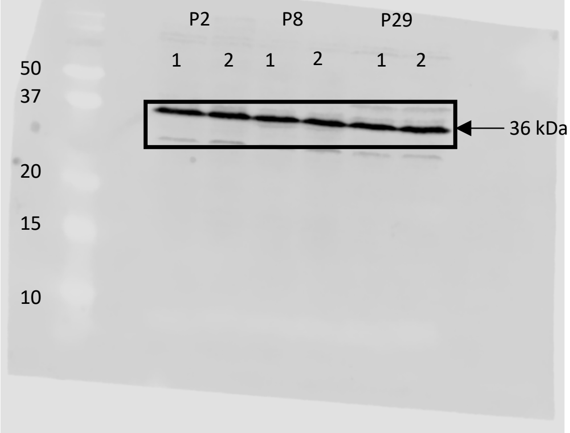


#### Set 2

phospho 4EBP1-S65

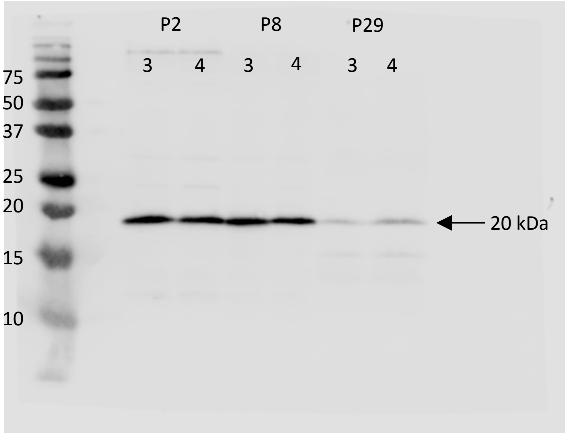


Total 4EBP1

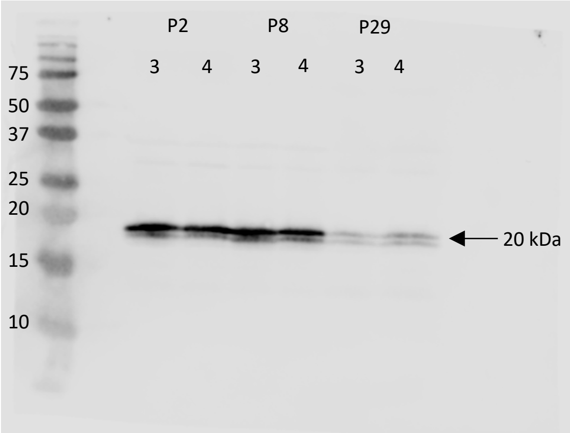


GAPDH

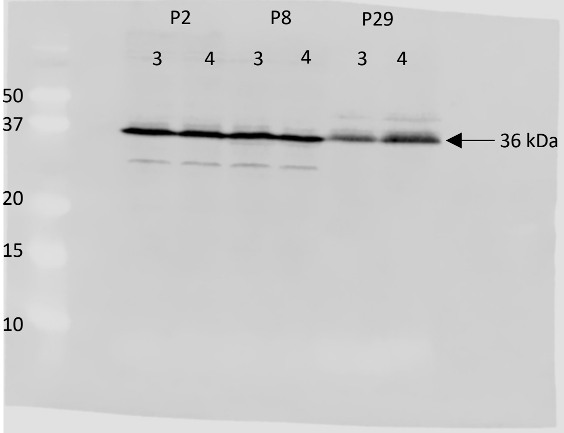

### Figure 1D: Regenerative

#### Set 1

phospho 4EBP1-S65

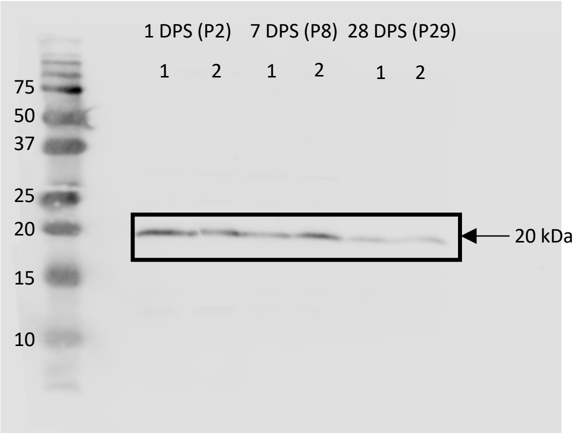


Total 4EBP1

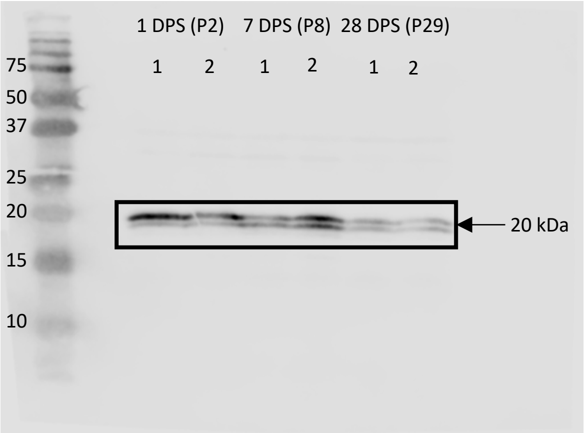


GAPDH

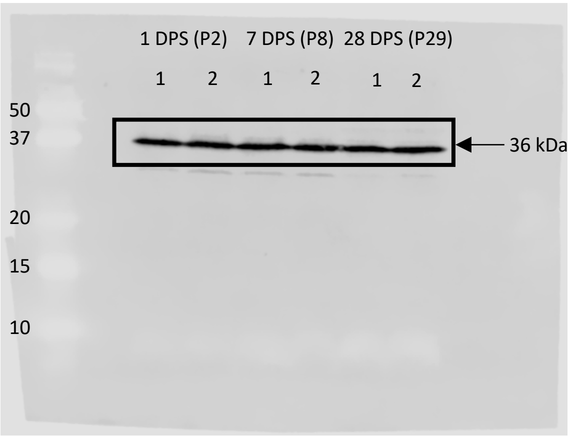


#### Set 2

phospho 4EBP1-S65

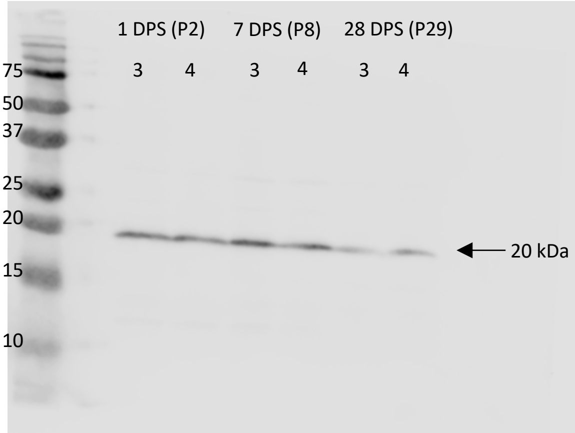


Total 4EBP1

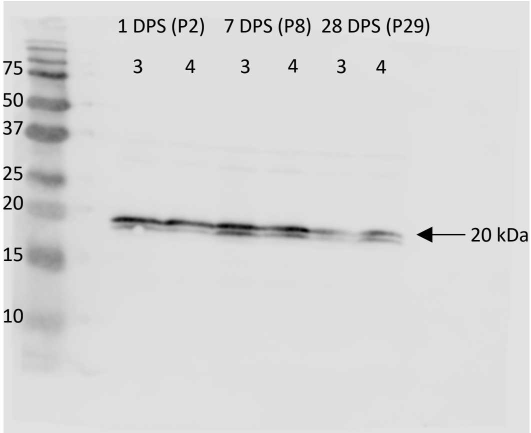


GAPDH

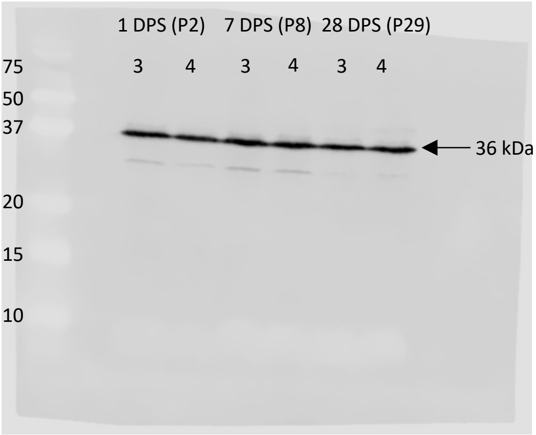

### Figure 1D: Non-Regenerative

#### Set 1

phospho 4EBP1-S65

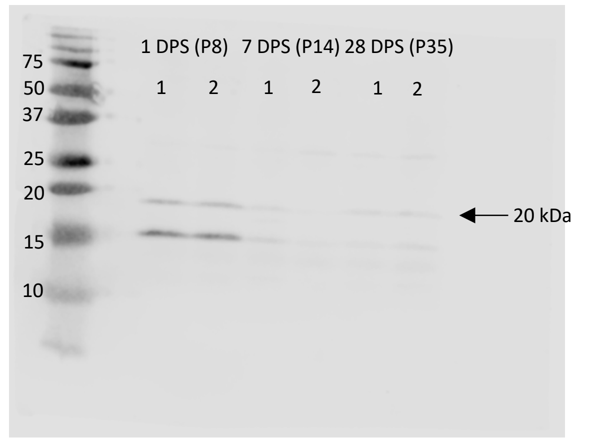


Total 4EBP1

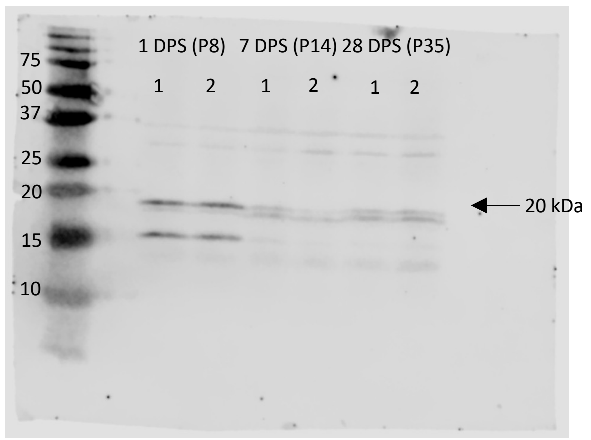


GAPDH

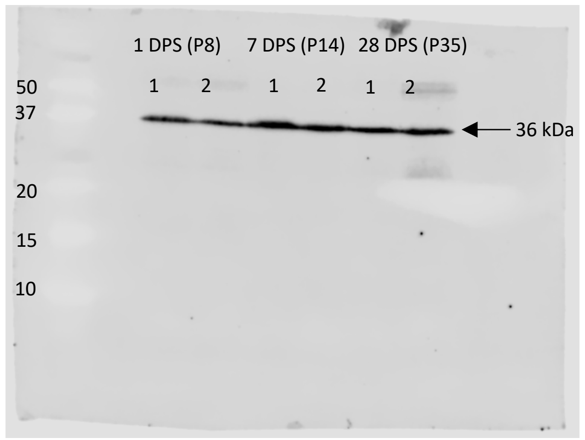


#### Set 2

phospho 4EBP1-S65

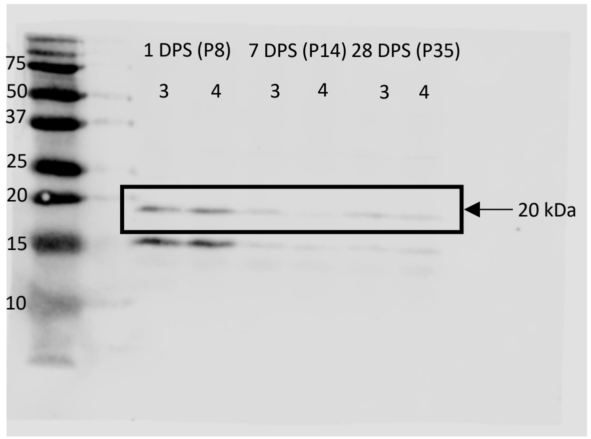


Total 4EBP1

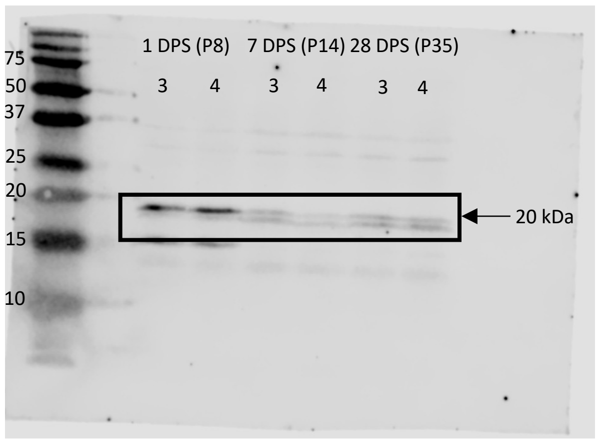


GAPDH

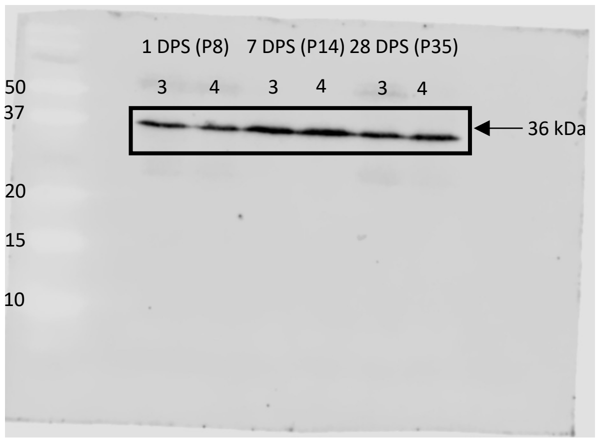

### Supplemental Figure 2A

Puromycin

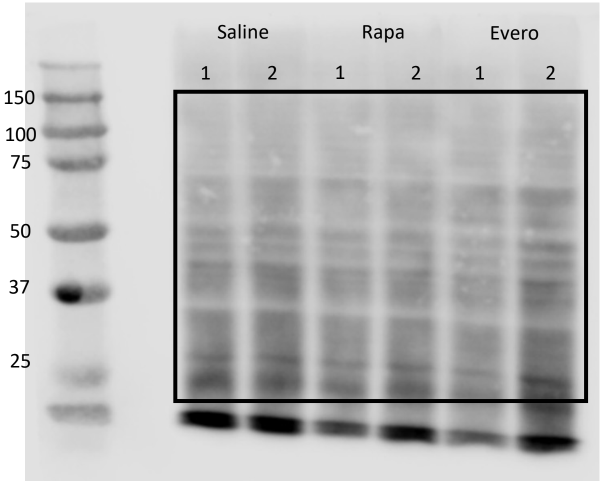


Puromycin

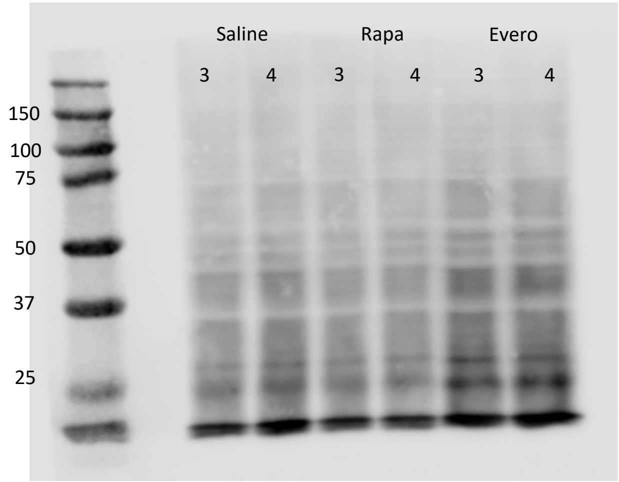


Ponceau Stain

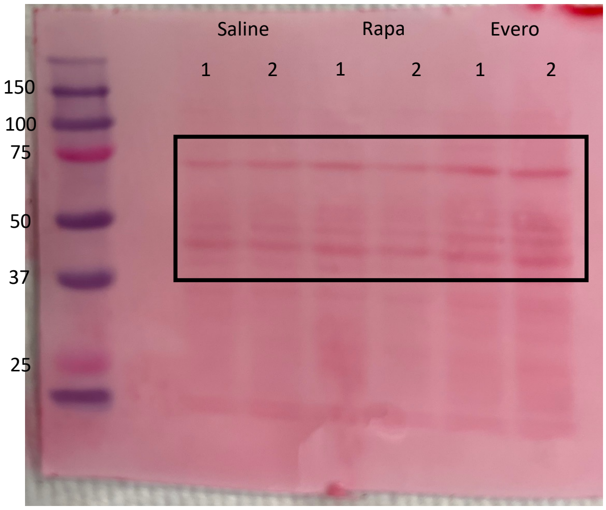


Ponceau Stain

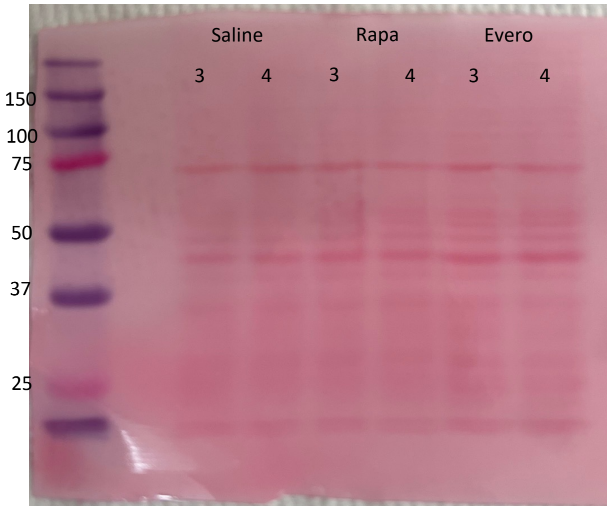

### Supplemental Figure 2B

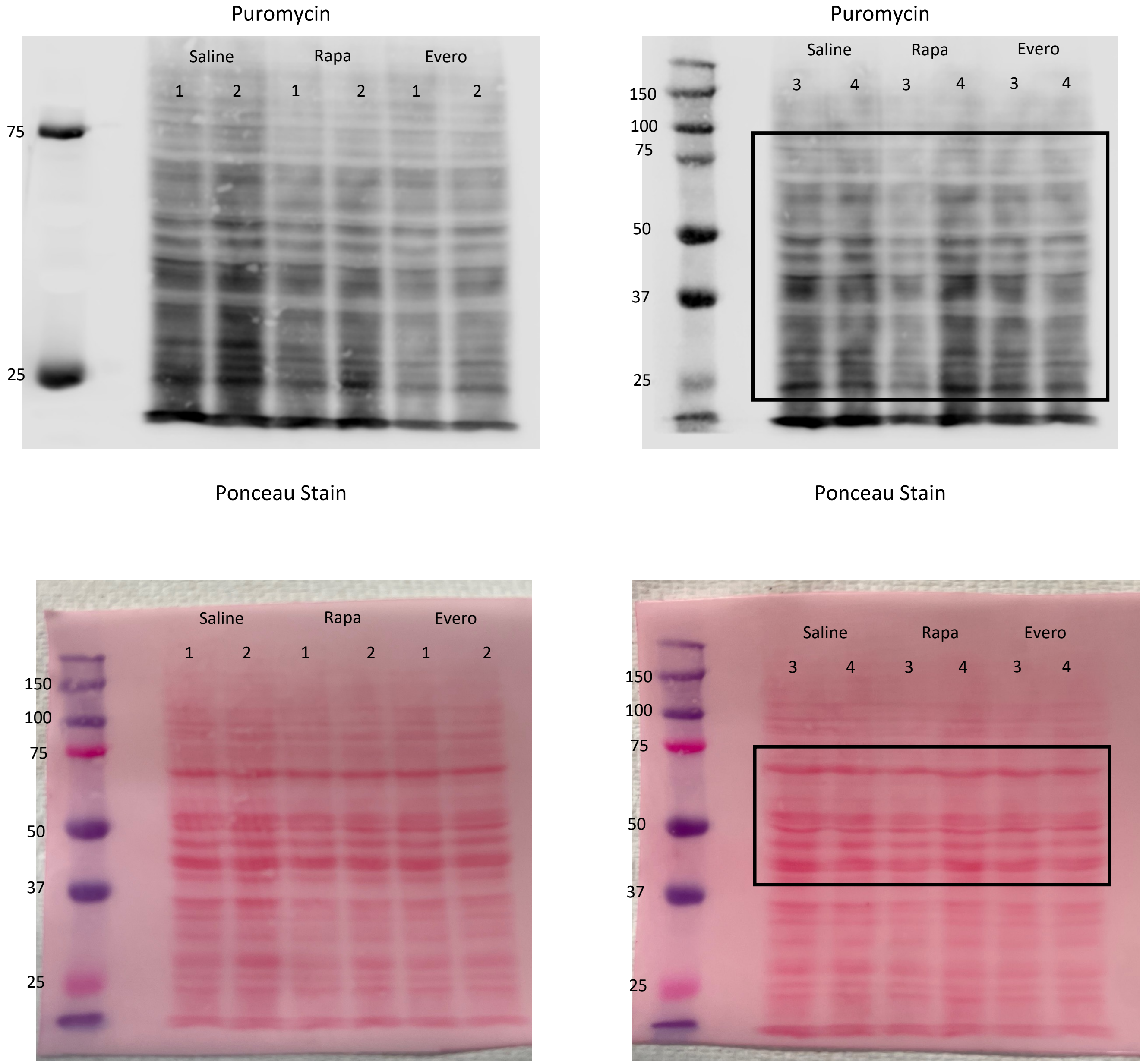

### Supplemental Figure 3A

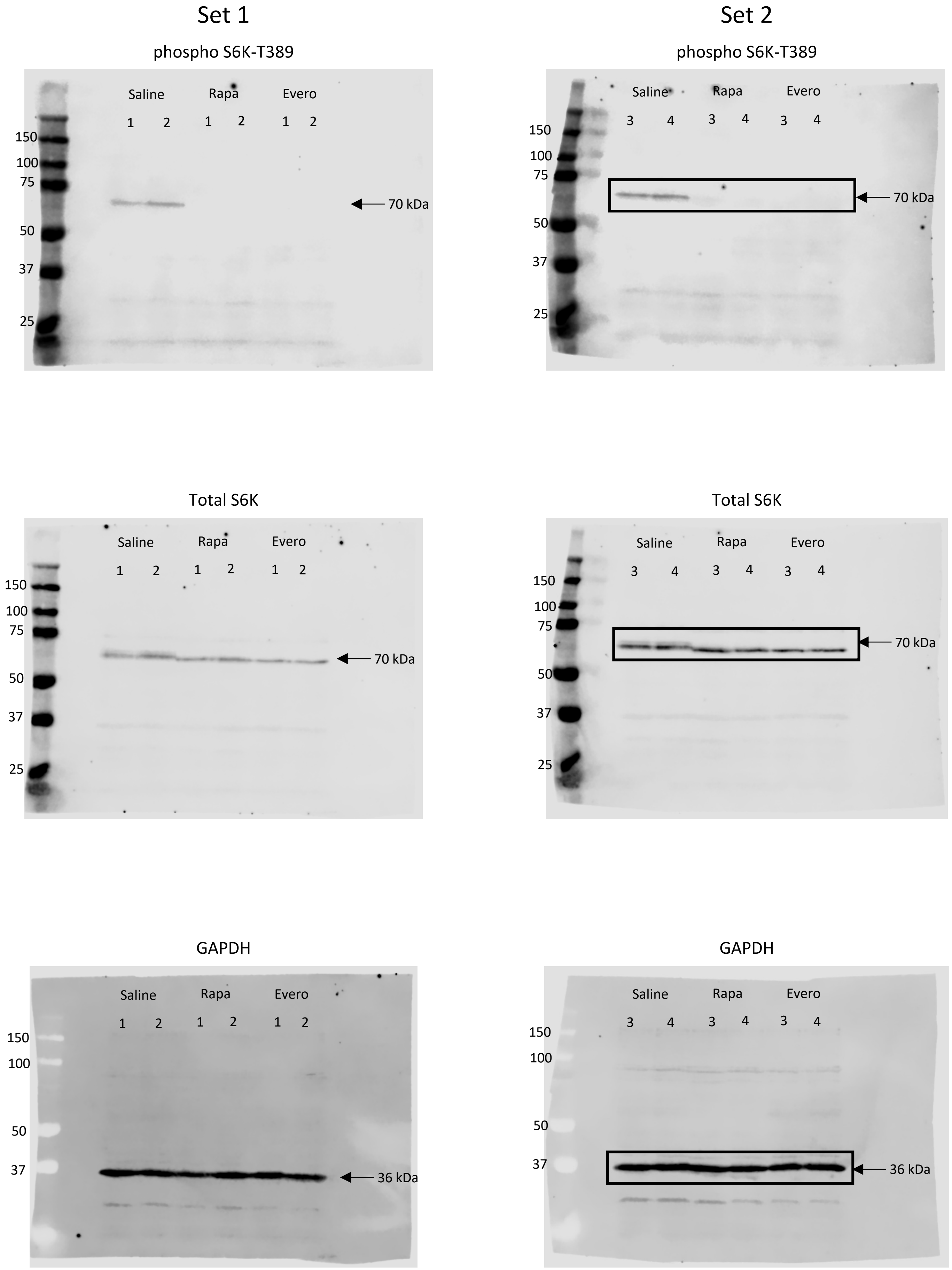

### Supplemental Figure 3B

Set 1

phospho 4EBP1-T37/46

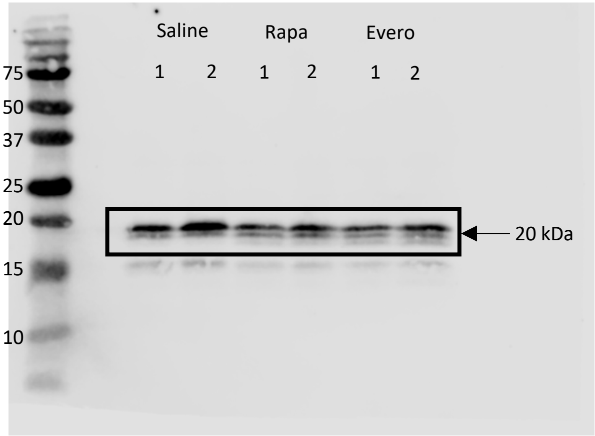


Set 2

phospho 4EBP1-T37/46

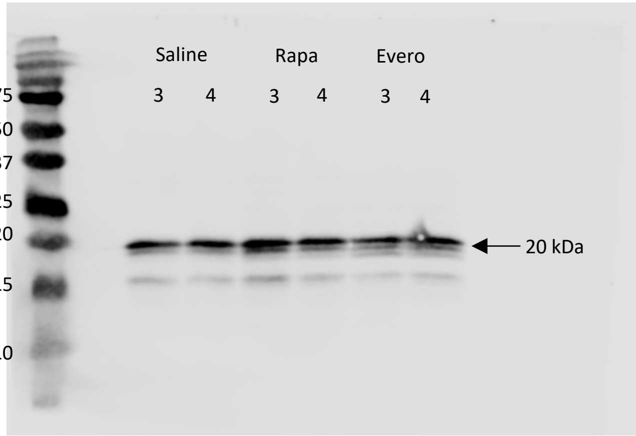


Total 4EBP1

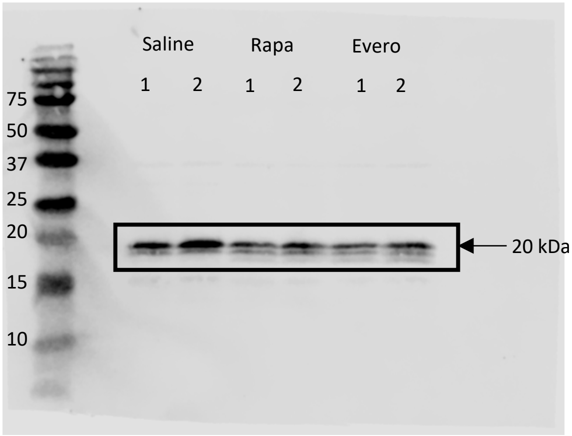


Total 4EBP1

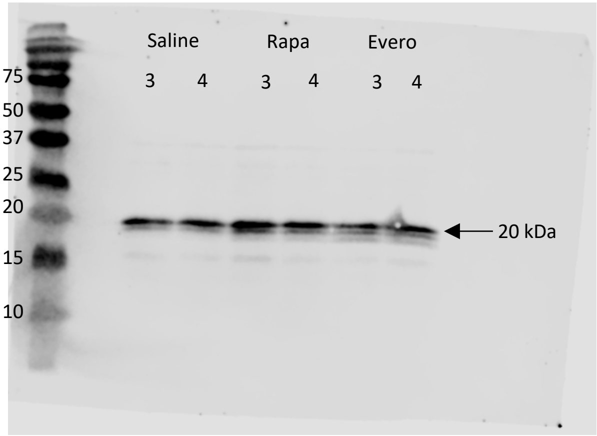


GAPDH

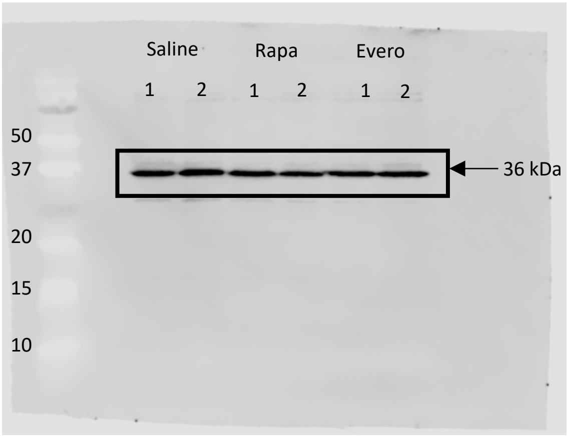


GAPDH

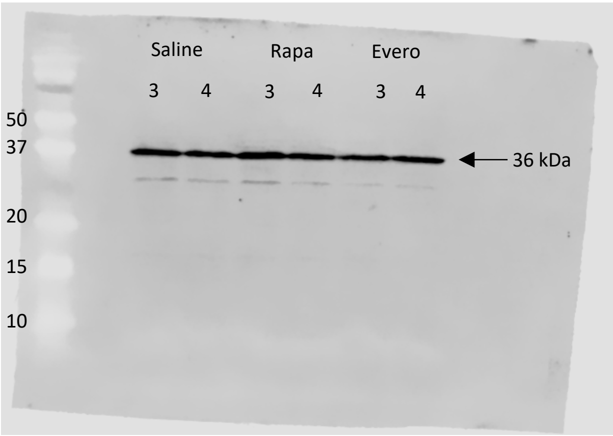

### Supplemental Figure 3C

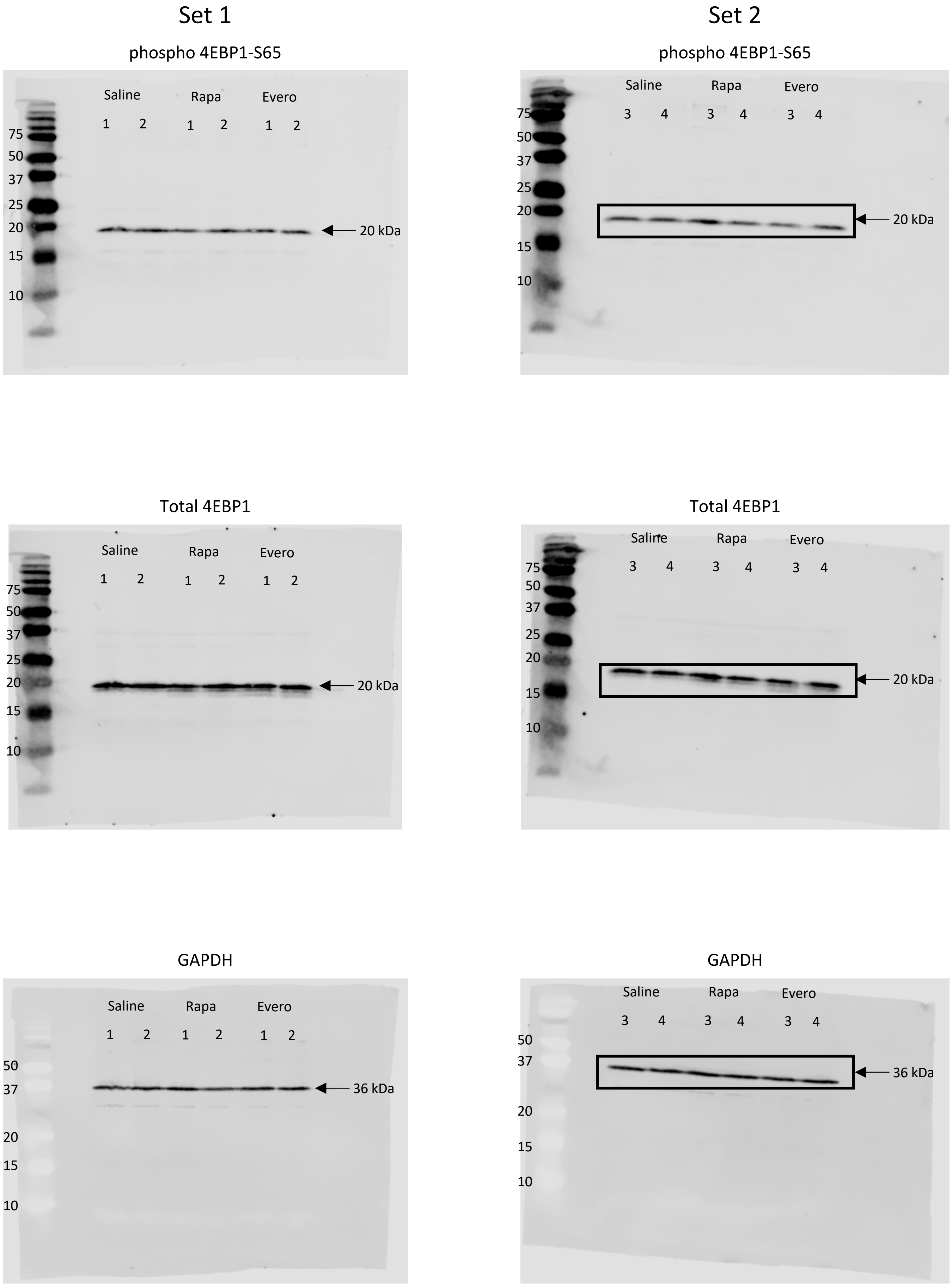

### Supplemental Figure 4A

#### Set 1

phospho S6K-T389

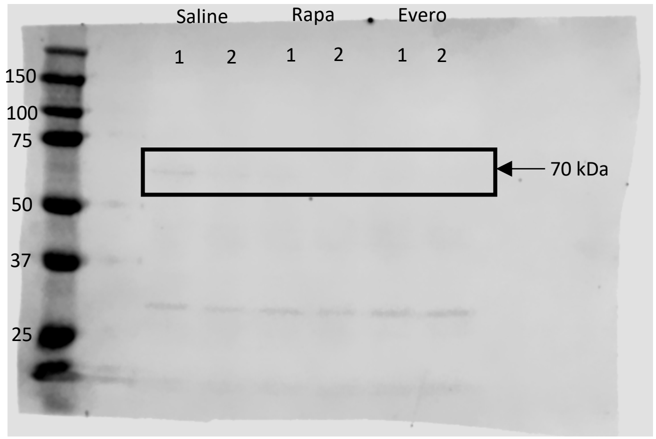


Total S6K

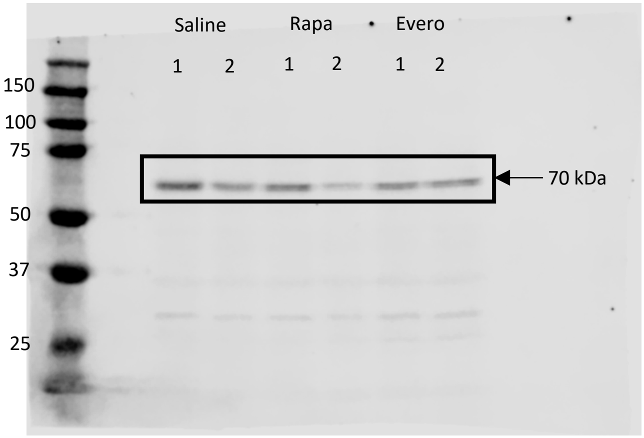


GAPDH

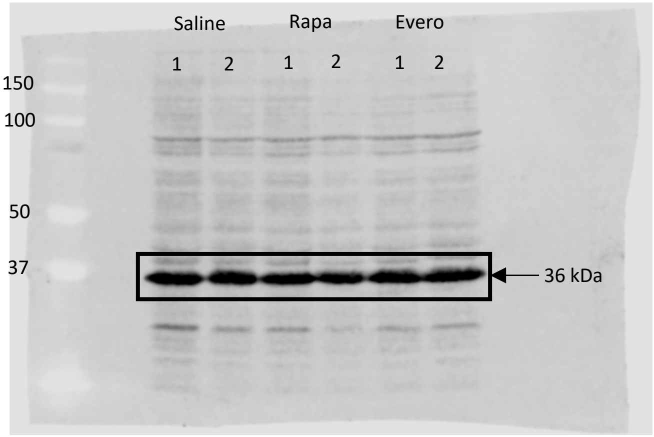


#### Set 2

phospho S6K-T389

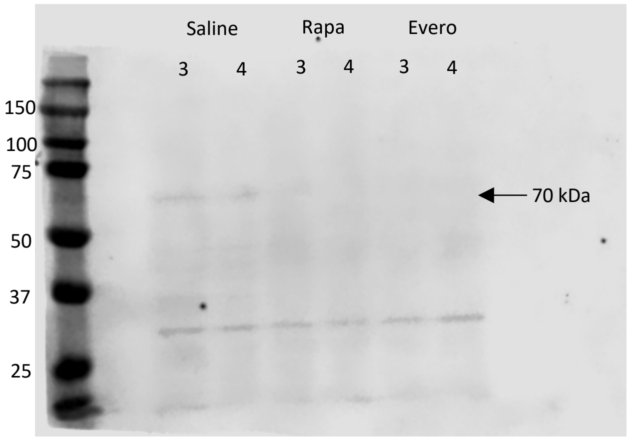


Total S6K

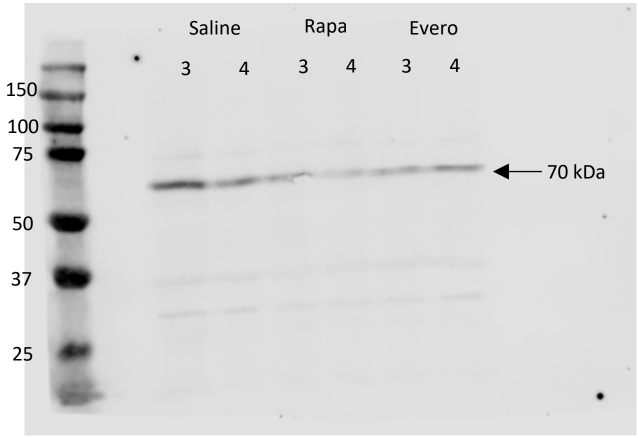


GAPDH

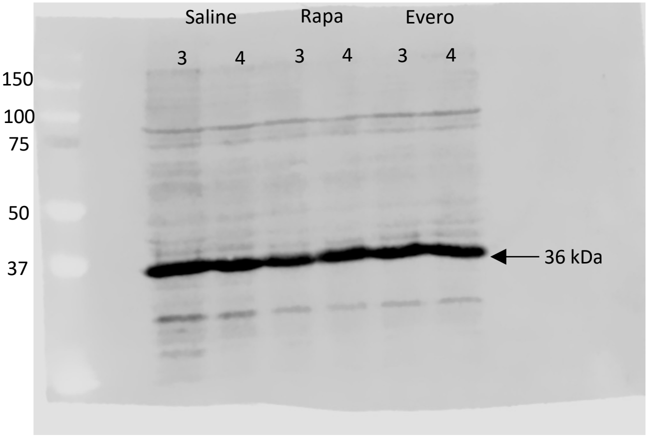

### Supplemental Figure 4B

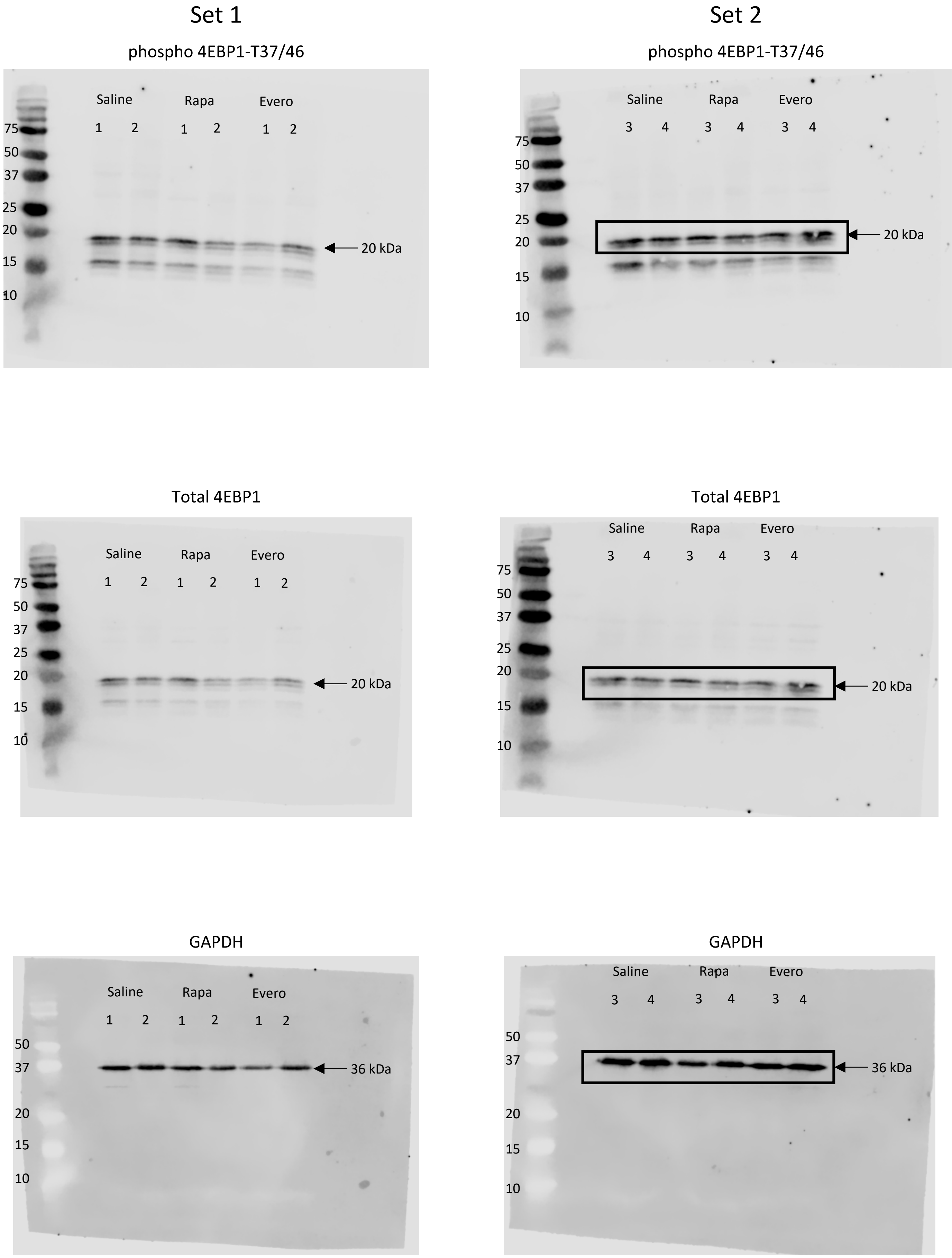

### Supplemental Figure 4C

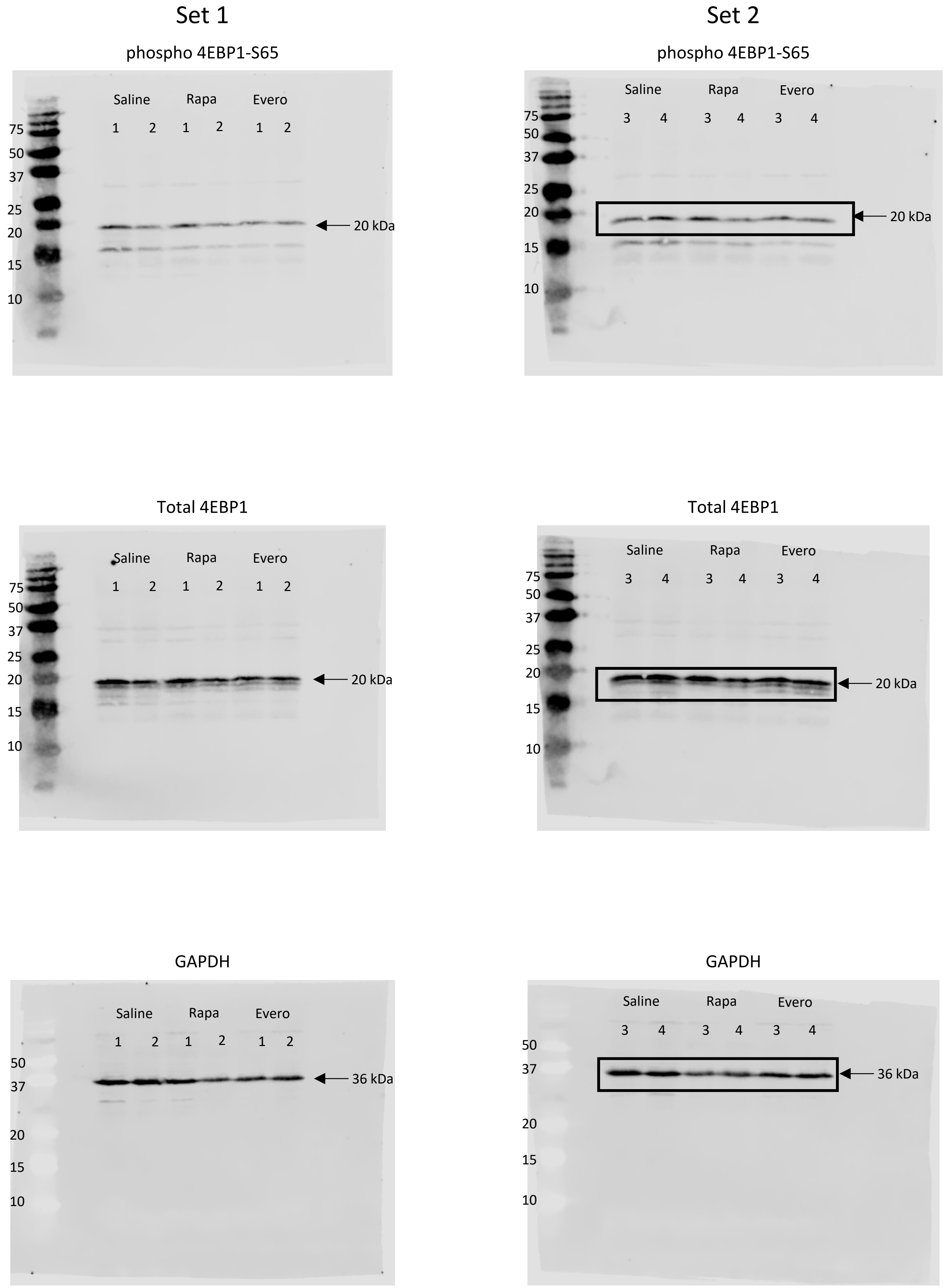
